# Downregulation of *Trpv4* and *Klf2* in brain microvessels is associated with the progression of neurovascular dysfunction and cognitive impairment in a model of heart failure with preserved ejection fraction

**DOI:** 10.1101/2025.01.08.631937

**Authors:** Sara M.P. Lambrichts, Laura van der Taelen, Irene Pastor, Peter Leenders, Nicole Bitsch, Daria Majcher, Denise Hermes, Steven J. Simmonds, Marcel van Herwijnen, Rick Kamps, Ellen Weltjens, Hellen P. Steinbusch, Floor Arentz, Maximilian Wiesmann, Amanda J. Kiliaan, Florian Caiment, Martina Kutmon, Elizabeth A.V. Jones, Robert J. van Oostenbrugge, Sébastien Foulquier

**Affiliations:** Dept of Pharmacology and Toxicology, Maastricht University, The Netherlands; Dept of Neurology, Maastricht University Medical Center, The Netherlands; Dept of Psychiatry and Neuropsychology, Maastricht University, The Netherlands; Dept of Cardiovascular Science, Centre for Molecular and Vascular Biology, KU Leuven, Belgium; Maastricht Centre for Systems Biology and Bioinformatics (MaCSBio), Maastricht University, The Netherlands; Dept of Translational genomics, Maastricht University, The Netherlands; Department of Medical Imaging, Anatomy, Research Institute for Medical Innovation, Radboud University Medical Center, Donders Institute for Brain, Cognition & Behaviour, Preclinical Imaging Center PRIME, Radboud Alzheimer Center, Nijmegen, the Netherlands; GROW – Research institute for Oncology and Reproduction, Maastricht University, The Netherlands; MHeNS – Research institute of Mental Health and Neuroscience, Maastricht University, The Netherlands; CARIM – Research institute of Cardiovascular Diseases, Maastricht University, The Netherlands

**Keywords:** Diastolic dysfunction, Cerebral small vessel disease, Microvascular dysfunction, Cerebrovascular reactivity, *Trpv4*, *Klf2*

## Abstract

Vascular cognitive impairment (VCI) shares major risk factors with heart failure with preserved ejection fraction (HFpEF), including obesity, diabetes and hypertension. Yet VCI research often relies on single-stimulus models, whereas patients experience combined risk factors. We therefore assessed cerebrovascular and cognitive phenotypes in an HFpEF model and investigated underlying mechanisms.

Male Lean and Obese ZSF1 rats underwent longitudinal assessments of blood pressure, glucose, cardiac function, and behavioural performance. Cerebral blood flow and neurovascular coupling were assessed by laser speckle contrast imaging. White matter integrity, blood–brain barrier (BBB) permeability, and vascular density were analysed by (immuno)histochemistry. Cortical microvessels were isolated for transcriptomic profiling, and selected targets were validated using multiplex in-situ hybridization.

Obese rats exhibited neurovascular uncoupling and impaired short- and long-term memory and spatial learning, accompanied by brain atrophy and reduced myelin. BBB permeability increased at 22-23 weeks and vascular density at 34-35 weeks in Obese vs Lean rats. Transcriptomic analysis of brain microvessels revealed altered processes related to angiogenesis, vasoreactivity, immune mechanisms and vascular remodelling, with consistent downregulation of *Trpv4* and *Klf2*.

Obese ZSF1 rats develop progressive neurovascular dysfunction associated with HFpEF onset and reduced *Trpv4* and *Klf2* expression in cerebral microvessels, two key vasoprotective genes.

## INTRODUCTION

From all dementia cases, more than 20% of the patients are diagnosed with vascular cognitive impairment (VCI). This mental disease covers all cognitive disorders caused by vascular dysfunction ranging from mild cognitive impairment to frank dementia (1). The most common cause of VCI is cerebral small vessel disease (cSVD), an umbrella term that covers all pathologies affecting cerebral microvessels and of which age and cardiovascular risk factor related cSVD is the most common form. This later type is characterized by white matter lesions, cerebral microbleeds, enlarged perivascular spaces, lacunar infarct, and brain atrophy (2). A systematic review and meta-analysis demonstrated that the presence of lacunes, white matter lesions and cerebral microbleeds are associated with left ventricular hypertrophy, highlighting a link between heart and brain disorders (3). Interestingly, the development of both VCI and heart failure are strongly linked to the presence of comorbidities such as obesity, diabetes, aging and hypertension (4). The brain and heart are especially sensitive to the chronic effect of comorbidities because of their high energetic and metabolic demands (5). With a growing aging population and the high prevalence of vascular risk factors in elderly, the incidence of VCI and heart failure is expected to further rise in the coming decades (6, 7).

Current evidence indicates that cognitive function declines more rapidly in patients with heart failure and that it may be affected prior to heart failure diagnosis (8). Two types of heart failure can be distinguished: heart failure with reduced ejection fraction (HFrEF), which is linked to the loss of cardiomyocytes caused by for instance ischemia or myocarditis; and heart failure with preserved ejection fraction (HFpEF), which is preceded by chronic comorbidities resulting in left ventricular hypertrophy and diastolic dysfunction (9).

Microvascular dysfunction has been proposed as a potential common mechanism for the development of cardiac and cognitive disorders (5). Although this process manifests differently in each organ, microvascular dysfunction can generally be defined as an altered structure of the microvasculature, such as vascular remodelling or rarefaction; and/or an altered function, including increased microvascular resistance and deregulation of vascular tone in reaction to stimuli, ultimately leading to hypoperfusion of the affected organs (10). Recent findings demonstrate that the coronary flow reserve is impaired in 50% of patients that present with coronary microvascular dysfunction (11). Moreover, the integrity of the microvascular wall barrier was found to be affected in HFpEF-associated comorbidities, thereby contributing to increased permeability of the endothelium. In case of cSVD patients, blood-brain barrier (BBB) impairment and cerebral hypoperfusion have been found to be strongly correlated with white matter lesions (12). Both cerebral blood flow and the BBB are controlled by cells of the neurovascular unit (13). To ensure the high energetic and metabolic demand of the brain, proper neurovascular coupling is essential. However, when this process becomes impaired, the metabolic demands of neurons cannot be fulfilled, subsequently leading to neurodegeneration and cognitive decline (2).

At present, no targeted therapeutic options are available for VCI nor HFpEF. Life quality of patients can only be improved by treating the associated comorbidities (6, 14). Although the underlying pathological mechanism of both disorders remains elusive, a central role for chronic hypoperfusion or impaired vascular reactivity through endothelial dysfunction has been proposed (5, 11). Moreover, recent findings demonstrated a reduced oxygenation reserve in myocardial and cerebral vasculature of patients with heart failure (5). Taken together, studies are highlighting microvascular dysfunction as an essential element of VCI in heart failure, but the mechanisms underlying this dysfunction remain largely unknown.

In this study, we aimed to characterize the cognitive and cerebrovascular phenotype in a rat model of HFpEF and to investigate the molecular processes governing the development of VCI in HFpEF. We performed a longitudinal study to assess the effect of comorbidities (obesity, diabetes, hypertension) on cerebral blood flow, neurovascular coupling, vascular density and BBB integrity at key time points associated with the disease progression. Furthermore, we assessed the dynamic transcriptomic regulations within cortical brain microvessels and validated *Trpv4* and *Klf2* as important signalling elements for the development of VCI in this HFpEF model.

## METHODS

### 1. Animals and study design

All experiments and procedures within this study were conducted in accordance with institutional and ARRIVE guidelines and approved by the local (Maastricht University IvD, DEC-UM) and national (CCD) ethical Committees for Animal Experiments (AVD10700202010326). ZSF1 rats and Wistar rats were obtained from Charles River Inc. and Janvier Labs, respectively. Obese ZSF1 rats are known to develop hypertension, obesity, diabetes and heart failure with preserved ejection fraction (HFpEF), whereas Lean animals present with only hypertension (15). The cognitive and cardiovascular phenotype of the ZSF1 rats was studied at an early time point before the development of HFpEF (14-16w in Study 1, 6-7w in Study 2, a middle time point (20-22w) where Obese ZSF1 present with HFpEF, and a late time point of prolonged exposure to comorbidities (30-32w) (16). In addition, Wistar rats (20-22w) were included in Study 2 to compare parameters such as cerebral blood flow measurements to reference values from normotensive rats.

Animals were socially housed in a 12/12-h reversed light/dark cycle (lights on from 19:00 to 7:00h), except during the novel objection recognition task (NORT, section 0). Cage locations within the housing rooms were randomized at the start of the experiments and at regular intervals during the study to minimize location bias. A radio played softly, providing background noise 24h a day, also during testing. Rats had free access to sterilized tap water and food; breeding diet (v1124-703, SSNIFF Spezialdiäten GmbH) for ZSF1 rats and standard diet (v1534-703, SSNIFF Spezialdiäten GmbH) for Wistar rats. Body weights were measured weekly. Due to the obvious difference in size/body weight between Obese and Lean animals, the investigators were aware of the group allocation during the execution of the studies. However, data sets were anonymized for all image analyses.

#### Study 1

Lean and Obese ZSF1 rats (n_Lean_=15, n_Obese_=14) were subjected to a series of behavioural testing to assess vascular cognitive impairment (VCI) at 14-16, 20-22 and 30-32 weeks old. Every two weeks, blood was collected via the *vena saphena* and fasted blood glucose (FBG) was measured (section 2). At 33-34 weeks old, thinned skull window surgeries (section 5) were performed to measure cerebral blood flow and assess neurovascular coupling using laser speckle contrast imaging (LSCI, PeriCam PSI HR, Perimed). At time of sacrifice, blood was collected from the *vena cava*. Isolated rat brains were cut sagitally and prepared for cortical brain microvessel isolation (section 10.1) and immunohistochemistry (section 0) respectively.

#### Study 2

The cardiovascular phenotype of Lean and Obese ZSF1 rats (n=14/group) was assessed in a longitudinal study during which blood pressure and echocardiography measurements were performed at the age of 6-7w, 20-21w and 30-31w. Every five weeks, blood was collected and FBG was measured. Thinned skull window surgeries and LSCI were performed on Lean and Obese rats at 6-7, 20-21 and 32-33 weeks old, as well as on normotensive Wistar rats (n=12, 20-21w). At time of sacrifice, blood was collected via *vena cava*. After isolation, rat brains were sagitally cut and used for cortical brain microvessel isolation (section 10.1) and immunohistochemistry (section 0) respectively.

### 2. Glucose measurement

Animals were fasted for 6h in cages with paper bedding. Blood was collected from the *vena saphena* using EDTA-precoated tubes (Microvette CB 300, Fisher Scientific, 16.444.100) during the active phase of the animals (14:00 to 16:00h). Briefly, glucose was measured using a Contour glucose meter (Ascensia Diabetes) and Contour test strips (Ascensia Diabetes, 84638099). Blood samples were kept on ice until the end of the procedure, followed by centrifugation at 2000 g for 10 minutes at 4°C. Plasma was then stored at −80°C until further use.

### 3. Assessment of the cardiovascular phenotype

Blood pressure and cardiac function were determined in Lean and Obese ZSF1 rats (n=14/group) at three disease stages. Systolic and diastolic blood pressures were measured in awake rats during their active phase with the CODA tail-cuff system (Kent Scientific corporation) using volume pressure recording. Prior to start of the experiment, animals were habituated to a restrainer for 15 minutes in a warming cabinet (28°C) for three consecutive days. At least 20 continuous blood pressure measurements were recorded and averaged for each rat.

During echocardiography, rats were anaesthetized with an i.p. injection of ketamine (75-100 mg/kg, Ketamine 10%, Alfasan) and xylazine (7.5-10 mg/kg, Xylasan 2%, Alfasan) dissolved in 0.9% NaCl (Baxter). All measurements were conducted according to previously described methods in Cuijpers et al. 2020 (16). During measurement, animals were placed in supine position and body temperature was monitored using a rectal probe and maintained at 37°C using a heating path and lamp. Heart and breathing rate were monitored via echocardiogram recording. A MS250 transducer (12.5-45 MHz) connected to Vevo 2100 echocardiography (VisualSonics) was used for 2D B- and M-mode imaging, pulse wave and tissue Doppler imaging. Left ventricle wall thickness was measured in B- and M-mode of parasternal short axis at the level of papillary muscles. Pulse wave Doppler measurements were performed to acquire pulmonary and aortic peak velocities in apical four-chamber view. Lastly, diastolic dysfunction was assessed by pulse wave and tissue Doppler imaging of the mitral valve. After the procedure, anaesthetized animals were awakened with atipamezole (i.p., 0.1 mg/kg). Vevo 2100 1.6.0 Software (VisualSonics) was used to extract functional parameters and perform analyses. Values of three stable cardiac cycles were averaged for each parameter.

### 4. Behavioural testing

#### Elevated zero maze

Anxiety of ZSF1 rats was assessed with the EZM in dark conditions. Rats were placed in the middle of an open arm to facing one of the closed arms and allowed to explore the maze over a period of five minutes. The total duration and distance travelled in the open and closed parts were measured via an infrared video camera connected to a video tracking system (Ethovision Pro, Noldus). Analysis was accomplished in Ethiovision XT15 (Ethovision Pro, Noldus).

#### Y-maze

The YM alternation task was performed to investigate spatial working memory in similar conditions as the EZM. Animals were placed in one of the three arms and allowed to explore the maze over a period of six minutes. The number of arm entries per animal was counted, and the alternation rate was used as a measure for working memory. Animals with a proper working memory have an alternation rate significantly above the 50% chance level. In addition, the total distance travelled in the YM was measured via an infrared video camera connected to a video tracking system and analysed in Ethiovision XT15. Animals with less than five arm entries were excluded from the dataset.

#### Novel object recognition task

Short-term memory of ZSF1 rats was evaluated using the NORT. Animals were housed individually two days prior to testing until the final testing day of the NORT (17). The test was performed in dark conditions and animals were familiarized to the arena and objects during two training days. On the two testing days, a first trial (T1) was performed with two identical objects. After a 1h-interval, animals were subjected to a second trial (T2) where one of the two objects was replaced by a novel object. Both trials lasted three minutes and animals returned to their home cage after the first trial. Total exploration time was monitored during testing: exploration of the objects by the animals was represented by looking, licking, sniffing or touching the objects while sniffing. Any rat exploring less than five and eight seconds during T2 of the first and second testing day respectively was excluded from the study. The discrimination index d2 during the first minute of the test was used as a measure for short-term spatial memory and is expressed as the difference between the time spent to the new and old object in T2 divided by the total exploration time in T2. Proper short-term memory is considered when d2 is significantly above 0.

#### Barnes maze

Assessment of long-term spatial learning and memory was done during a five-day testing period in an adapted BM task (18). The experiment was performed under light conditions and a maximum of six rats was allowed to habituate to the testing environment for 15 minutes prior to the start. During the four training days, ZSF1 rats were subjected to four trials a day with each rat being allocated to a different target arm. The trial ended after three minutes or when the animal entered the correct target arm. On the last day of the BM, a probe trial was performed in which the animal remained in the arena for three minutes. The relative time spent in the target quadrant during the first 30 seconds of the trial was used as a measurement for long-term memory. Lastly, a reversed learning trial was performed in which each rat was assigned to a different arm than during the training days to assess reversal learning. The total duration and distance travelled until reaching the correct target arm was measured with a video camera connected to a video tracking system. Analysis was done in Ethiovision XT15.

### 5. Thinned skull window surgeries and laser speckle contrast imaging

Animals underwent thinned skull window surgeries to assess cerebral blood flow and neurovascular coupling. One hour following carprofen injection (s.c., 5 mg/kg, Rimadyl), rats were anaesthetized using 2.5-3.5% isoflurane (IsoFlo, Zoetis) and fixed in a stereotactic frame. During surgery, body temperature was monitored using a rectal probe and maintained at 37°C using a heating path and lamp. Before midline incision, 2% lidocaine (Fresenius Kabi AG) was injected onto the periosteum. The region above the left barrel cortex (5.5-6 mm lateral and 2.5 mm posterior to Bregma) was thinned using a high-speed microdrill (NSK Euope, Mio 35M) under a stereo microscope (Leica MZ8, Leica Microsystems). After completing the skull-thinning, animals were injected with 0.05-1 mg/kg medetomidine (s.c., SedaSTART, ASTfarma) to maintain a light sedation state. To keep the skull hydrated and improve image quality, light mineral oil (0.838 g/ml, Sigma Aldrich, 330779) and heavy mineral oil (0.862 g/ml, Sigma Aldrich, 330760) were subsequently added on top of the thinned window. After a 20-minute isoflurane washout period, whisker stimulation was performed by mechanically shaking the whiskers for 30 seconds at a frequency of 5 Hz on the contralateral side of the thinned skull window. The stimulation was done using a home-made stepper-motor system connected to a foam brush and controlled by an Arduino board. Stimulation was repeated three times with a two-minute interval and images were acquired at baseline before and during stimulation via LSCI (resolution = 10 μm, image size = 5 x 5 mm; frame averaging = 10; frame rate = 44 images/second). After measurements, animals were s.c. injected with saline and atipamezole (Antisedan, Orion Corporation) to end sedation. All rats received carprofen (5 mg/kg) in their drinking water and were checked daily until three days post-surgery.

Analysis was accomplished using PIMSoft 1.11 software (Perimed); three regions of interest (0.1 x 0.1 mm) were selected in the barrel cortex area. Neurovascular coupling was relatively expressed to the average baseline cerebral blood flow prior to each whisker stimulation. Three values were averaged per animal.

### 6. Sample collection

One hour prior to sacrifice, rats were injected with 0.05 mg/kg buprenorphine (s.c., Bupaq, Fendigo, Pharma AG). Briefly, animals were anaesthetized with 4% isoflurane and blood was withdrawn from the *vena cava*, followed by pentobarbital injection (i.v., 100 mg/kg, Nembutal, Ceva) and cardiac perfusion with ice-cold PBS supplemented with 1% EDTA (pH 8.0, ThermoFisher, 15575020). Organs, plasma and urine samples were prepared for further analyses and either snap frozen in liquid nitrogen or fixed in 4% paraformaldehyde (PFA, Sigma-Aldrich, 158127), washed with PBS and stored in PBS with 0.1% sodium azide (Sigma Aldrich, 26628-22-8). In addition, brains and hearts were weighted on a precision scale (Mettler Toledo, XP603S), whereas tibia length was measured using a vein caliper.

### 7. Plasma molecular analysis

During terminal procedure, blood was collected via *vena cava* from non-fasted animals in EDTA-precoated tubes (BD Vacutainer K2E, 367864, BD) and centrifuged at 2000 g for 10 minutes. Plasma was stored at −80°C until further use. Triglyceride levels were determined by the clinical chemistry department (Centraal Diagnostisch Laboratorium, MUMC).

### 8. Immunohistochemistry

Brains were sliced into free-floating sections of 30 μm thickness using a vibratome (Leica VT1200, Leica Microsystems). For each immunofluorescent staining, two brain slices (Bregma +1.6 mm; Bregma +2.2 mm) were included and two FOVs per section were imaged from deep cortical regions. Coronal sections were stored in PBS with 0.1% sodium azide until staining. First, slices were washed with PBS supplemented with 0.1% Triton-100 (PBS-T, Sigma-Aldrich, X100), PBS and PBS-T for 10 minutes and blocked with 1% bovine serum albumin (BSA, VWR, 422361V) diluted in PBS-T for 1.5h at room temperature. Briefly, sections were stained overnight (4°C) with anti-rat IgG-biotin (1:500 dilution in PBS-T, Jackson ImmunoResearch, 712-065-150) to assess blood-brain barrier (BBB) permeability, followed by incubation with streptavidin Alexa Fluor^TM^ 594 conjugate (ThermoFisher, S11227) for 2h. Following washing steps, cerebral vessels were stained using a tomato lectin (1:100 dilution in PBS-T, Lycopersicon Esculentum (tomato) Lectin, DyLight 649, Vector Laboratories, DL-1178). Finally, sections were mounted with antifading mounting medium (Invitrogen, P36980).

#### Vascular density

Image stacks (x=275 μm, y=275 μm, z=10 μm, 2 μm step size) of lectin-stained brain sections (n=4-10/group) were acquired on a confocal microscope (Leica DMI 4000B, Leica Microsystems) at a 40x magnification using Leica Application Suite Advanced Fluorescence (LAS AF) software (Leica Microsystems). For each FOV, a maximal intensity projection was prepared in ImageJ (FIJI Distribution, NIH). Vascular density, branching points and average vessel length were determined with Angiotool (19). The following parameters were applied: vessel diameter: 3-8; intensity: 50-255; remove small particles: 40. Images were excluded from the analysis when the signal-to-noise ratio was incompatible with the Angiotool analysis. For each animal, images from two brain slices were averaged. The analysis was performed by an experimenter blinded to the groups.

#### BBB permeability

To identify the presence of IgG leakages (n=8-10/group), brain sections were scanned using an ImageXpress Pico Automated Live Cell Imager (Molecular Devices, San Jose, USA) at 10x magnification (1 pixel = 0.69 μm). Acquired images were automatically stitched by the software (CellReporterXpress software, Molecular Devices). BBB permeability was determined by the extravasation of IgG from vessels into the parenchymal tissue. The number and size (surface area) of the leakages were assessed in Arivis Vision 4D (version 3.6.2, Zeiss). In short, a machine learning segmentation was performed to identify background (tissue negative for lectin and IgG signals), vessels (lectin-positive structures) and IgG proteins (IgG-positive signal). A subtraction was performed to identify “IgG objects” outside the “vessels objects” i.e. extravascular IgG signal. The obtained IgG leakages were then filtered to remove artificial IgG background signal (objects <500 µm^2^) and the number and size (surface area, µm^2^) of IgG leakages was exported for analysis.

### 9. Myelin integrity

A Luxol fast blue staining of brain sections from ZSF1 and Wistar rats was performed to estimate white matter area and myelin content of the pericallosal white matter (defined as the corpus callosum and adjacent lateral white-matter tracts, including the external capsule). The protocol was adapted from Santiago et al. 2018 (20). Free-floating sections (Bregma +2.2 mm) were transferred to adhesive glass slides to be air-dried overnight. After 24h incubation with Luxol fast blue (Solvent Blue38, Carl Roth GmbH, 7709.3), differentiation of the white matter was completed using 0.1% lithium carbonate (Merck, 105676). After washing steps with 70% ethanol (Boom B.V., 64-17-5) and demi water, sections were incubated with cresyl violet (Carl Roth GmbH, 7651.2) followed by differentiation in 6% acetic acid-ethanol (Acetic Acid, ThermoFisher, S25118A) for five minutes each. Finally, sections were mounted with Entellan (Sigma-Aldrich, 1.07961) and imaged at 10x magnification using a BX50 microscope (Olympus) connected to a MBF camera (MBF Bioscience CX9000) and a MAC5000 Controller system (Ludl Electronic Products Ltd, 73 05020). Images were visualized and processed with

Stereo Investigator software (version 2020, MBF Bioscience, MicroBrightField). The Cavalieri method was used to estimate the pericallosal white matter and whole brain area. The ratio of the pericallosal white matter area over whole brain area was then used to assess white matter atrophy. Myelin content was assessed using LFB mean gray value (MGV) intensities. LFB images were converted from ScanScope Virtual Slide files to TIFF files using Qupath (Qupath-0.5.1., Belfast, Northern Ireland). An in-house built ImageJ script was developed to perform background corrections. The background corrected LFB images were processed using ImageJ. The ROIs covering the pericallosal white matter (corpus callosum and external capsule) and the striatum were delineated using a custom ImageJ script. During this analysis, LFB images were converted to a grayscale format, from which MGV data was extracted. To increase interpretability, LFB values were inverted and converted into percentages. Decreased relative LFB corresponds to decreased myelin content.

In addition, myelin integrity was assessed using polarized light imaging (PLI). PLI is a novel imaging technique that leverages the birefringent properties of myelin sheaths to assess fiber orientation in histological brain sections (21–24). By transmitting polarized light, this method enables quantitative estimations of fiber orientation and inclination within a given sample (22). PLI has the potential to identify early indicators of WM damage, surpassing the diagnostic sensitivity of traditional histochemical techniques such as LFB staining (21). The sections were mounted onto glass microscope slides and dried at 37 °C overnight. Subsequently, dried slides were coated with polyvinylpyrrolidone (PVP; Sigma-Aldrich, 9003-39-8, Darmstadt, Germany) and covered with 24×60 mm coverslip glasses in a laminar flow cabinet. To ensure optimal preservation, the prepared slides were left to dry under controlled conditions for two weeks. The slides were then imaged using a PLI microscope (Zeiss Axioplan microscope, Germany) equipped with a RGB-camera (Axiocam ERc 5s, Zeiss, Germany). Background correction images were taken at the start of every imaging session to be utilized during post-processing. Bregma +0.48 mm was selected from the atlas of the rat brain to be imaged for this study. Two FOVs were acquired; each FOV was acquired in nine sequential images at rotation between 0 and 160 degrees. The raw images underwent post-processing in Matlab (MATLAB R2020b; MathWorks Inc., Natick, MA, USA), wherein images were fitted to Jones Calculus. Background corrections were performed and FOVs were stitched together for each of the nine sequential images per rat brain, facilitating the construction of PLI maps; dispersion, FOM-HSV, inclination, in-plane, retardance, and transmittance. The retardance (myelin density), dispersion (myelin quality), and inclination maps were then imported into ImageJ to delineate the pericallosal white matter (corpus callosum and external capsule) and the striatum as ROIs using a Wacom IntuosPro Small pen tablet (Model PTH-451, Düsseldorf, Germany). MGV was recorded for dispersion and retardance for each ROI per section. For retardance, the higher the MGV, the better the myelin density. In standard interpretation, lower MGV scores for dispersion are indicative of better myelin quality. However, to enhance interpretability, dispersion values were inverted and both the inverted dispersion data and the untouched retardance MGV values were converted into percentages. Within this framework, higher relative dispersion and retardance values correspond to improved myelin quality and density.

### 10. Gene expression analysis of brain microvessels

#### Isolation of brain microvessels

Cortical brain microvessels were freshly isolated from one cerebral hemisphere immediately after sacrifice. The cortex was homogenised in cold, sterile PBS using a tissue grinder and centrifuged for five minutes at 2000 g. The resulting pellet was then suspended in 15% dextran-PBS (Sigma Aldrich, 31390) and centrifuged at 4300 g for 15 minutes. After dissolving the pellet in PBS containing 1% BSA (Fatty Acid-free, Sigma Aldrich, 126609), the solution was transferred to a 100 μm cell strainer (Pluriselect, 43-57100-51), followed by filtration through a 20 μm nylon mesh (Pluriselect, 43-50020-03). Brain microvessels remaining on the latter filter were collected using 0.5% BSA-PBS (Fatty Acid-free, Sigma Aldrich, 126609) and centrifuged at 4300 g for 10 minutes. The final pellet was either suspended in DPBS and air-dried for immunohistochemistry, or TRIzol (ThermoFisher Scientific, 15596018) for RNA isolation, and stored at −80°C.

#### Characterization of brain microvessels

Microvessels were washed and permeabilized with 0.1% IGEPAL (Sigma Aldrich, CA-630) in TRIS-buffered saline (TBS), TBS and 0.1% IGEPAL-TBS for 5 minutes. Unspecific binding was blocked with 3% donkey serum in TBS for 1h. Briefly, primary antibodies were diluted in 0.1% IGEPAL-TBS supplemented with 0.3% donkey serum and incubated overnight (4°C): goat anti-pericyte-derived growth factor receptor β (PDGFR-β, 1:100, R&D systems) and rabbit anti-ionized calcium-binding adapter molecule 1 (Iba-1, 1:500, FUJIFILM Wako Chemicals, 019-19741). Following washing steps, samples were incubated for 1h at room temperature with corresponding secondary antibodies dissolved in 0.1% IGEPAL-TBS supplemented with 0.3% donkey serum: donkey anti-goat AF488 (1:200, Invitrogen, A-21202) and donkey anti-rabbit AF488 (1:500). To detect astrocytic endfeet, samples were first blocked with 1% BSA + 3% donkey serum in 0.1% IGEPAL-TBS, followed by overnight incubation with rabbit anti-aquaporin 4 (1:1000, Sigma, A5971) and visualized with donkey anti-rabbit AF488 (1:500) diluted in 0.1% IGEPAL-TBS containing 1% BSA. Following washing steps, cerebral microvessels were stained overnight with tomato lectin diluted in 0.1% IGEPAL-TBS supplemented with 0.3% donkey serum at 4°C. Counterstaining of the nuclei was performed using NucBlue (Invitrogen, R37606) according to the manufacturer’s instructions. Finally, slides were mounted using antifading mounting medium.

#### RNA isolation, library preparation and bulk RNA sequencing

For samples of the first time point, microvessels from two rats (8-9w) were pooled into one sample before RNA isolation because of the young age of the animals. RNA was extracted from brain microvessels of Lean and Obese ZSF1 rats using TRIzol according to the manufacturer’s protocol and stored at −80°C until further processing. RNA purity and integrity (RIN) were checked according to the manufacturer’s protocol using Nanodrop One (ThermoFisher Scientific) and Agilent 2100 Bioanalyzer system (Agilent Technologies) respectively. Purification of the mRNA (n=4/group/time point) and library preparation were performed according to the manufacturer’s protocol using the RiboNaut™ rRNA Depletion Kit (NETFLEX, NOVA-512963, PerkinElmer) and the Rapid Directional RNA-Seq Kit 2.0 (NETFLEX, NOVA-5198, PerkinElmer), respectively. Lastly, sequencing was performed on an S1 flow cell using the Novaseq 6000 sequencing system (NovaSeq 6000, Illumina, Inc).

#### Transcriptomic analysis

Raw sequencing data was trimmed using fastp (25). The R package ‘DESeq2’ was used to identify DEGs using a p-value (false discovery rate; FDR) below 0.05 and a fold change above 1.5 (26, 27). Gene Ontology enrichment analysis was performed to identify overrepresented biological processes in the list of DEGs using the clusterProfiler R-package (v. 4.12.6, (28). Protein-protein interaction networks of selected GO terms were created using the stringApp (v. 2.1.1, (29)) and visualized in Cytoscape (v3.10.1) (30) using the RCy3 R-package (v. 2.24.0, (31)).

### 11. RNAscope assay

RNAscope^TM^ Multiplex Fluorescent Detection kit v2 (Advanced Cell Diagnostics, ACD, Bio-Techne Gmbh, 323120) was performed according to the manufacturer’s instructions. Specific mRNA probes were designed against rat *Klf2* (111711-C1) and *Trpv4* (476661-C2) targets. Three-plex positive and negative control probe mixtures were included in the experiment (**Figure S10**). All baking steps were performed in the ACD Hybez II Hybridization Oven (ACD, 321710).

The multiplex detection of mRNAs was performed on paraffin-embedded ZSF1 brains of Lean and Obese animals at 22-23 and 34-35 weeks of age. After incubating the sections with protease for 30 min, the solution was replaced with fresh protease PLUS for another 10-min incubation. Following probe hybridization, brain sections were stored overnight in saline sodium citrate buffer. TSA Vivid Fluorophores 570 (channel C1, 323272, 1:1500) and 650 (channel C2, 323273, 1:1500) were used to visualize *Klf2* and *Trpv4* probes, respectively. Lycopersicon Esculentum Lectin (DyLight 488, Vector Laboratories, DL-1174-1) was used to detect blood vessels following RNAscope procedure.

Slides were imaged (x=175 μm, y=175 μm, z= 6 μm, 0.5 μm step size) with confocal microscopy at a 63x magnification (Leica SPE, Leica Microsystems B.V. Amsterdam). For each rat brain (n=4/group), three FOVs were acquired and analyzed in Arivis Vision 4D (version 3.6.2, Zeiss, Rostock, Germany). Machine learning segmentation was performed to identify background, lectin-positive signal, and *Klf2* and *Trpv4* transcripts. Next, the compartments feature was applied to identify only *Klf2* and *Trpv4* signal inside the vessel objects. From this procedure, several parameters about the target expression in the vascular network were extracted: number of transcripts and total intensity. All parameters were corrected for the total vessel volume.

### 12. Statistical analyses

Group size was calculated a priori for both animal studies (see above description of Study 1 and Study 2) using G*Power3.1. For study 1, the discrimination index d2 has been chosen as primary outcome parameter for group size calculation due to its high variability compared to other cognitive readout parameters: ANOVA with repeated measures, effect size f=0.25, α error = 0.05; power β = 0.80, and considering a possible 15% drop-out rate, resulting in n=15 animals/group. For study 2, the change in CBF has been chosen as primary outcome parameter for group size calculation as it was central to answer our research question: ANOVA, effect size f=0.90, α error = 0.05; power β = 0.80, and considering a possible 5% drop-out rate, resulting in min n=11 animals/group for the primary readout parameter. Considering required sample sizes for post-mortem analyses and to be consistent with the group size in Study 1, the group size in Study 2 was increased to n=14 animals/group.

Data are represented as mean ± SEM. All statistical analyses were performed with GraphPad Prism 10.0.1 software (San Diego, USA). Normality distribution was tested with the Shapiro-Wilk test. Two-way analysis of variance (ANOVA) was used to compare Lean and Obese groups over time, followed by Sidak’s post hoc testing with correction for multiple comparisons. One-sample t-tests were performed to determine whether: the alternation percentage in ZSF1 rats was different from 50% during the YM; the discrimination index d2 was significantly different from 0 during the NORT; and whether cerebral blood flow, neurovascular coupling, vascular density, the number of branching points, white matter content, and BBB integrity of ZSF1 rats differed from Wistar rats. *P*-values <0.05 were considered statistically significant.

## RESULTS

In this study, we used male Zucker fatty and spontaneously hypertensive (ZSF1) rats as a model for HFpEF and comorbidities. For cardiovascular and cognitive characterisation, only Lean and Obese ZSF1 animals were studied, while Wistar rats were included as a normotensive reference group for the *in vivo* assessment of microvascular function and post-mortem IHC analyses.

### 1. Cardiovascular phenotype: Obese ZSF1 rats develop HFpEF

Systolic and diastolic blood pressures were increased in all ZSF1 rats at each time point compared to normotensive values reported in a previous publication (dashed line; single t-test: p_systolic_<0.0001, **Figure 1A**; p_diastolic_<0.01, **Figure S1A**) (32). Moreover, Obese ZSF1 rats showed a significantly higher blood pressure at 20w compared to Lean animals (p_systolic_<0.01, p_diastolic_<0.01). The development of obesity and diabetes was monitored over time; an increase in body weight (p<0.0001, **Figure S1B**) and FBG (p<0.05, **Figure 1B**) was observed in Obese rats starting from 5 and 10 weeks of age, respectively. In addition, triglyceride levels were elevated in Obese vs Lean at 22-23w and 34-35w (p<0.0001, **Figure 1C**).

**Figure 1:**
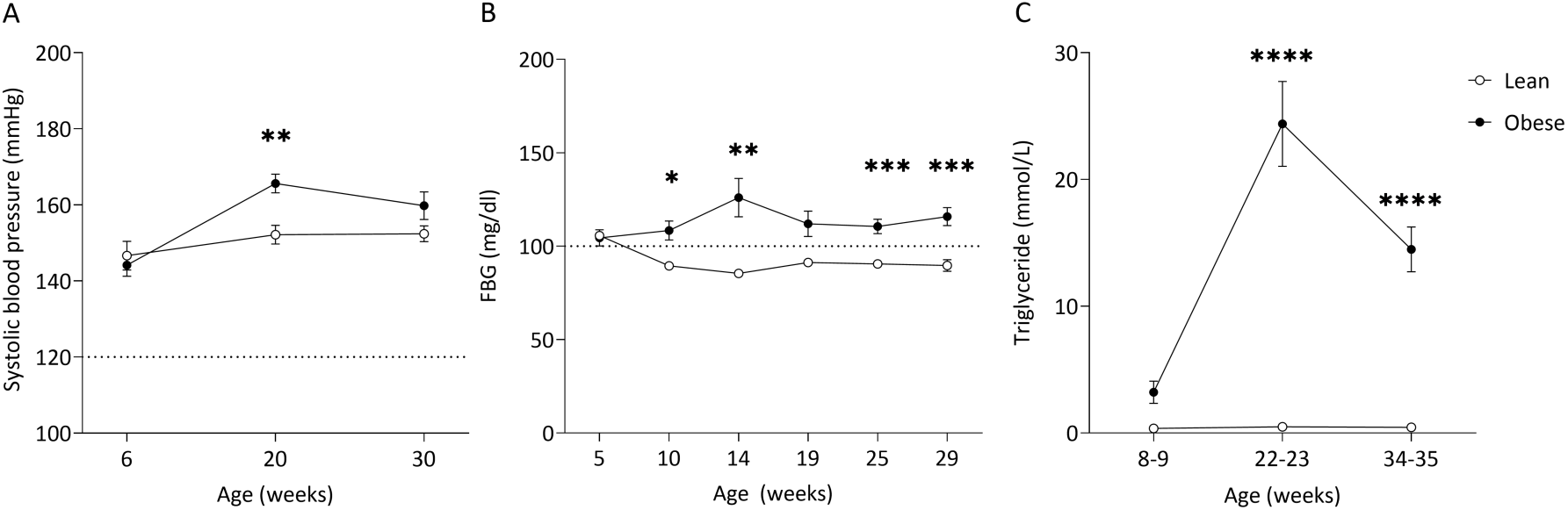
Progression of systolic blood pressure, blood glucose and triglyceride levels in ZSF1 Obese vs Lean rats. (A) Systolic blood pressure of Lean (open circles) and Obese (filled circles) ZSF1 rats (normotensive value t=120 mmHg, dashed line (32)): 2-way ANOVA (p_age_<0.0001, p_group_=0.01, p_age x group_=0.03). (B) Fasted blood glucose (FBG) levels (t=100 mg/dl; dashed line): mixed effects (p_age_=0.56, p_group_<0.0001, p_age x group_=0.0006). (C) Non-fasted blood triglycerides levels: 2-way ANOVA (p_age_<0.0001, p_group_<0.0001, p_age x group_<0.0001). Sidak’s multiple comparisons test: *p<0.05; **p<0.01; ***p<0.001 ****p<0.0001. n=14-15/group.

The progression of HFpEF in ZSF1 rats was assessed at the age of 7, 21, and 31 weeks using echocardiography. All animals showed a preserved ejection fraction (EF) above 50%, moreover, Obese ZSF1 rats had a significantly higher EF at all disease stages compared to Lean animals (p_7w_<0.001, p_21w_<0.01, p_31w_<0.0001, **Figure S2**). Pulse wave Doppler measurements of the mitral valve indicated a decrease in the ratio of rapid filling (early mitral inflow peak velocity, E) over atrial contraction (late mitral inflow peak velocity, A) in Obese animals at 21 and 31w (p<0.0001, **Figure 2A**). The isovolumetric relaxation time (IVRT) was significantly higher in Obese vs Lean at 21w and a tendency towards increased IVRT was observed at 31w (p_21w_<0.05, p_31w_=0.10 **Figure 2B**). This was accompanied by an elevated deceleration time in Obese ZSF1 rats at 21w (p<0.001, **Figure 2C**).

**Figure 2:**
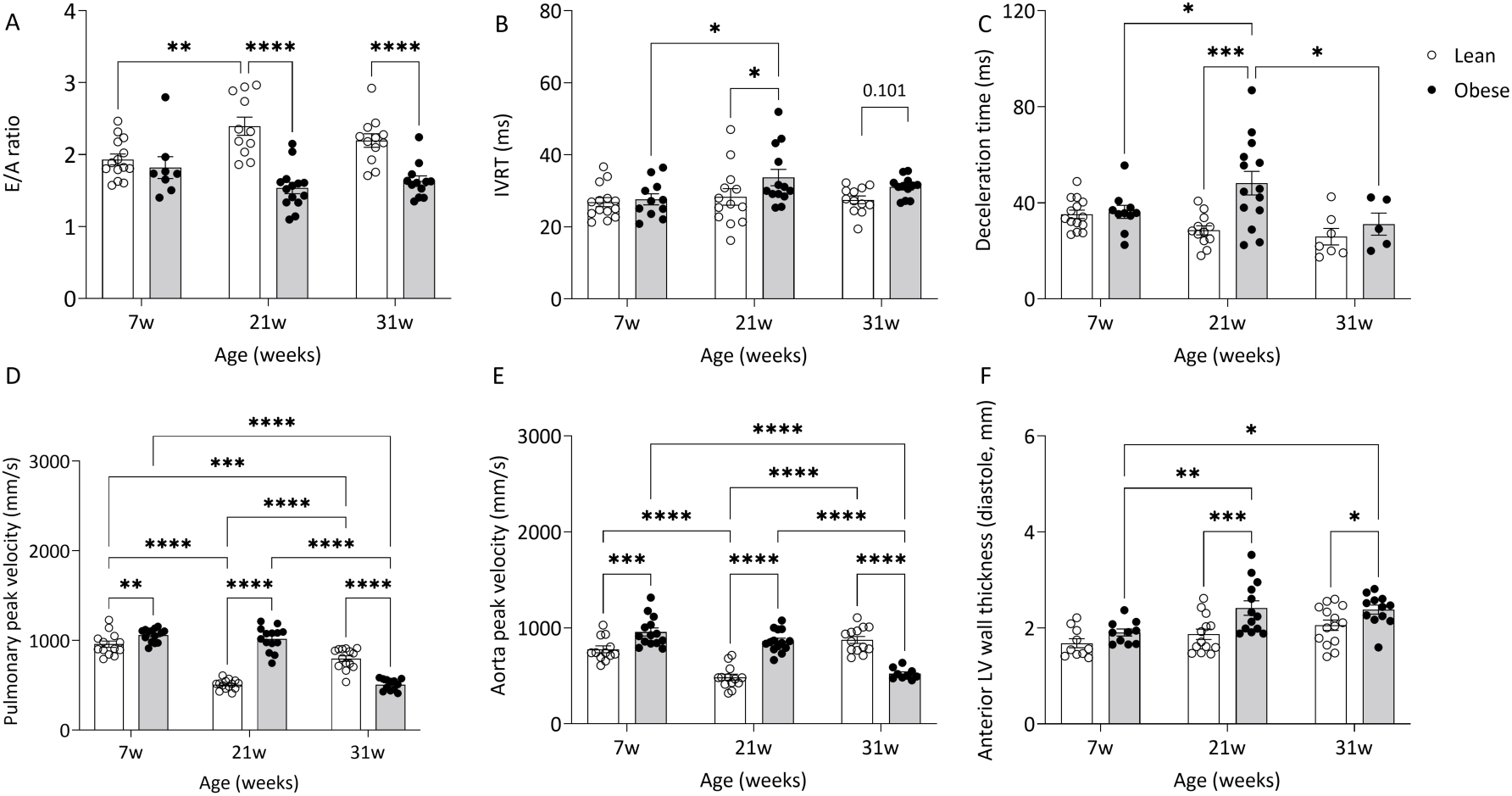
In vivo assessment of cardiac function in ZSF1 rats at different time points using echocardiography. (A) Ratio of early and late mitral inflow peak velocity (E/A ratio; mixed effects: p_age_=0.66, p_group_<0.0001, p_age x group_=0.0008), (B) isovolumetric relaxation time (IVRT mixed effects: p_age_=0.13, p_group_=0.013, p_age x group_=0.27), (C) deceleration time (mixed effects: p_age_=0.06, p_group_=0.0079, p_age x group_=0.0173), (D) pulmonary peak velocities (mixed effects: p_age_<0.0001, p_group_<0.0001, p_age x group_<0.0001), (E) aortic peak velocities (mixed effects: p_age_<0.0001, p_group_=0.0189, p_age x group_<0.0001), and (F) anterior left ventricular (LV) wall thickness (mixed effects: p_age_=0.0013, p_group_=0.0002, p_age x group_=0.35) in Lean (open circles; white bars) and Obese (filled circles; grey bars) ZSF1 rats. Sidak’s multiple comparisons test: *p<0.05, **p<0.01, ***p<0.001, ****p<0.0001. n=5-14/group.

Moreover, the measurement of pulmonary peak velocities (p_7w_<0.01, p_21w_<0.0001, **Figure 2D**) and aorta peak velocities (p_7w_<0.001, p_21w_<0.0001, **Figure 2E**) showed a significant increase in Obese vs Lean at the first two time points, while being decreased at 31w (p<0.0001). During diastole, anterior LV wall was thickened in Obese ZSF1 rats at 21w (p<0.001) and 31w (p<0.05). Other echocardiographic parameters are available in **Figure S2** and **Figure S3**. Altogether, the assessment of cardiac function confirmed HFpEF pathology in Obese ZSF1 rats at both 21w and 31w, and was furthermore associated with an elevated heart weight/tibia length ratio at each disease stage (**Figure S4**).

### 2. Cognitive function: impaired performance in Obese ZSF1 rats

A series of behavioural testing was performed to assess anxiety, working memory, short-term memory, and long-term spatial learning/memory using the elevated zero maze (EZM), Y-maze, novel object recognition task (NORT), Barnes maze (BM), respectively (**Figure 3A-D**). Obese ZSF1 rats were more anxious at 30-32 weeks old, spending less time in the open arms compared to Lean rats in the EZM (p<0.001, **Figure 3E**). Lean animals became less anxious over time (p_20-22w_<0.05 and p_30-32w_<0.01 vs 14-16w), whereas the number of open arm entries remained unchanged between groups (**Figure S5A**). During the YM task, working memory was not affected by comorbidities at any of the time points, only Lean rats showed an impaired working memory at 30-32w (single t-test: p<0.05 vs t=50 (33), **Figure 3F**). When comparing Obese rats of 30-32w to 14-16w, a significant decrease in working memory was observed. In addition, the percentage of alternations (p_age_<0.001) decreased over time, while the relative time spent in centre remained unchanged (**Figure S5C-D**). Lastly, locomotor activity in both the EZM and YM task was lower in Obese ZSF1 rats at 14-16w (**Figure S5B**). Although impaired working memory could not be confirmed in the Obese, Lean rats did have a significant impairment of working memory at the latest time point. This could be due to a decrease in the number of entries over time (p_age_<0.0001), especially in Obese animals at 20w and 30w of age. In addition, Obese rats were slightly more anxious than the Lean, which might have affected their capability to alternate as well during the YM task.

**Figure 3:**
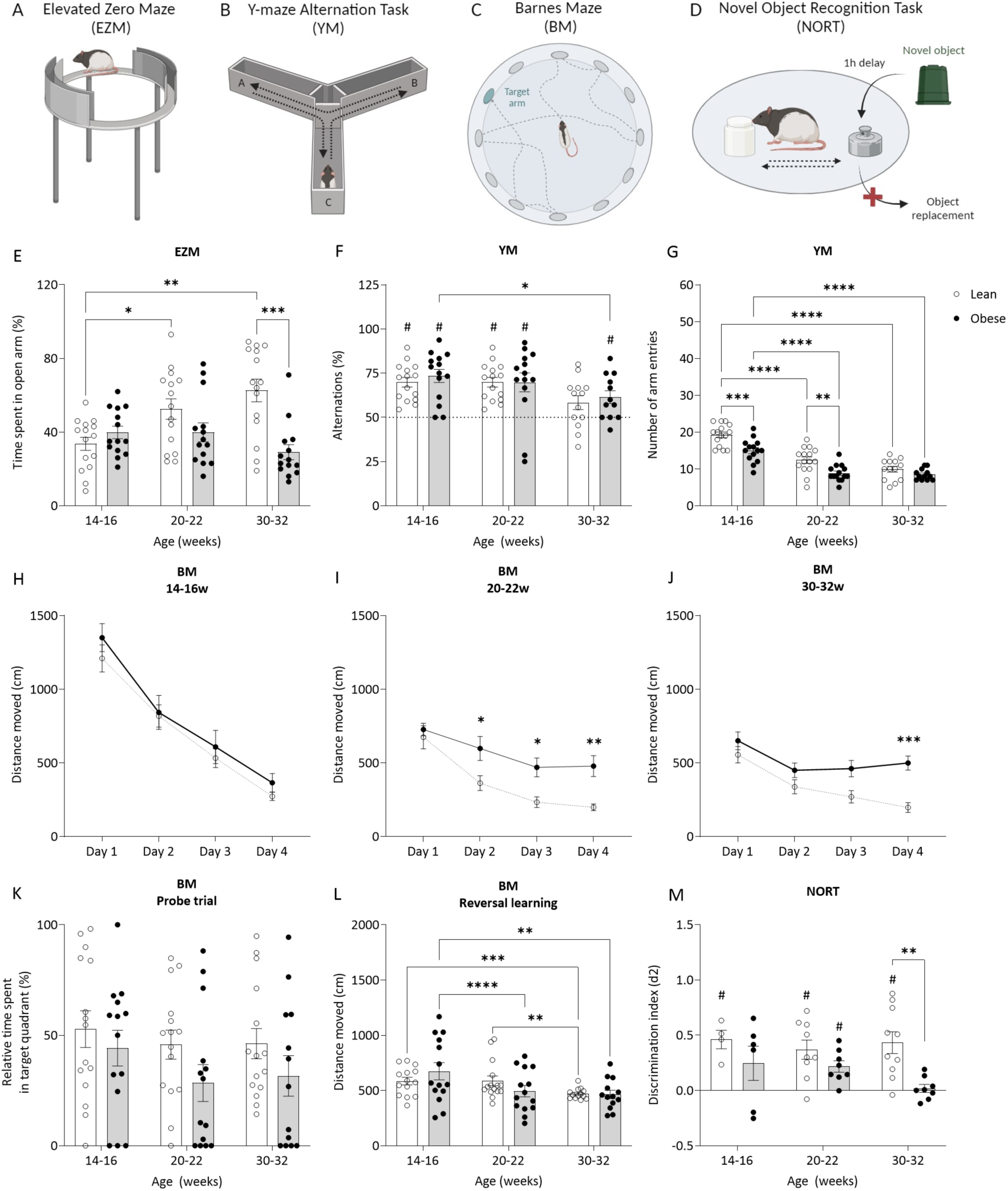
Anxiety and cognitive function of ZSF1 rats at three time points. Lean (open circles; white bars) and Obese (filled circles; grey bars) ZSF1 rats underwent a series of behavioural tests, including (A) elevated zero maze (EZM), (B) Y-maze alternation task (YM), (C) Barnes maze (BM) and (E) novel object recognition task (NORT) to assess anxiety, working memory, long term spatial learning/memory, and short-term memory, respectively. (E) EZM (2-way ANOVA: p_age_=0.12, p_group_=0.0006, p_age x group_=0.0012); (F) YM, percentage of alternations (mixed effects: p_age_=0.0019, p_group_=0.60, p_age x group_=0.83, (33)), (G) YM, number of entries (mixed effects: p_age_<0.0001, p_group_<0.0001, p_age x group_=0.15); (H-J) BM, spatial learning at 14-16w (2-way ANOVA: p_age_<0.0001, p_group_=0.46, p_age x group_=0.35), 20-22w (2-way ANOVA: p_age_<0.0001, p_group_=0.01, p_age x group_<0.0001), and 30-32w (mixed effects: p_age_<0.0001, p_group_=0.03, p_age x group_<0.0001), (K) BM 30-second probe trial, long-term memory (mixed effects: p_age_=0.30, p_group_=0.04, p_age x group_=0.84), (L) BM, reversal learning (mixed effects: p_age_<0.0001, p_group_=0.84, p_age x group_=0.01); (M) NORT (mixed effects: p_age_=0.43, p_group_=0.0019, p_age x group_=0.29); Sidak’s multiple comparisons test: *p<0.05, **p<0.01, ***p<0.001, ****p<0.0001. Single t-test: #p<0.05 vs (F) alternation rate of 50% (dotted line,) and (M) d2=0 (33). EZM: n=14-15/group, YM: 13-15/group; NORT: n=4-10/group; BM: 13-15/group. Illustrations in panels A-D were created using BioRender.com.

Two-way ANOVA revealed short-term memory impairment in Obese ZSF1 rats at 30-32 weeks of age (single t-test: p<0.05 vs d2=0) during the novel object NORT, and at this time point, the discrimination index d2 was significantly lower in Obese vs Lean (**Figure 3M**). Furthermore, total exploration time during this task was not different between groups (**Figure S5E**). Spatial learning was assessed in the BM task and was found to be impaired in Obese vs Lean starting from 20-22w (**Figure 3H-J**). In the 30-seconds probe trial, a significant decrease in long-term spatial memory was observed in Obese animals compared to Lean rats (p_group_<0.05, **Figure 3K**), and reversal learning was affected by aging (p_age_<0.0001, p_age x group_<0.01, **Figure 3L**). During this task, locomotor activity was significantly lower in Obese ZSF1 rats at all time points (**Figure S5F**).

### 3. Brain atrophy and early white matter deficits in Obese ZSF1 rats

Brain atrophy was observed in Obese vs Lean ZSF1 rats at 8-9w (p<0.0001) and 22-23w (p<0.0001, **Figure 4B**). Within Lean and Obese groups, brain size also decreased significantly over time (p<0.0001). The relative proportion of white matter area was also lower in Obese animals at 22-23w and 34-35w compared to Lean (p<0.05, **Figure 4C**). Moreover, within the Lean and Obese group, the relative size of the pericallosal area was significantly higher over time, due to the progression of brain atrophy. Myelin content, as measured by the relative LFB signal, was decreased in Obese vs Lean animals in both the pericallosal and striatum areas (pericallosum p_group_=0.04, **Figure 4D**; striatum p_group_=0.02, **Figure 4E**). The relative retardance, corresponding to myelin density, increased over time independently of the groups in both areas (pericallosum p_group_<0.0001, **Figure 4F**; striatum p_group_<0.001, **Figure 4G**). The relative dispersion, corresponding to myelin quality, was not affected by age in the pericallosal area but increased in the striatum area (p_age_<0.0001, **Figure 4I**). There was only a mild decrease observed between Obese and Lean groups in the striatum (p_group_=0,051, **Figure 4I**).

**Figure 4:**
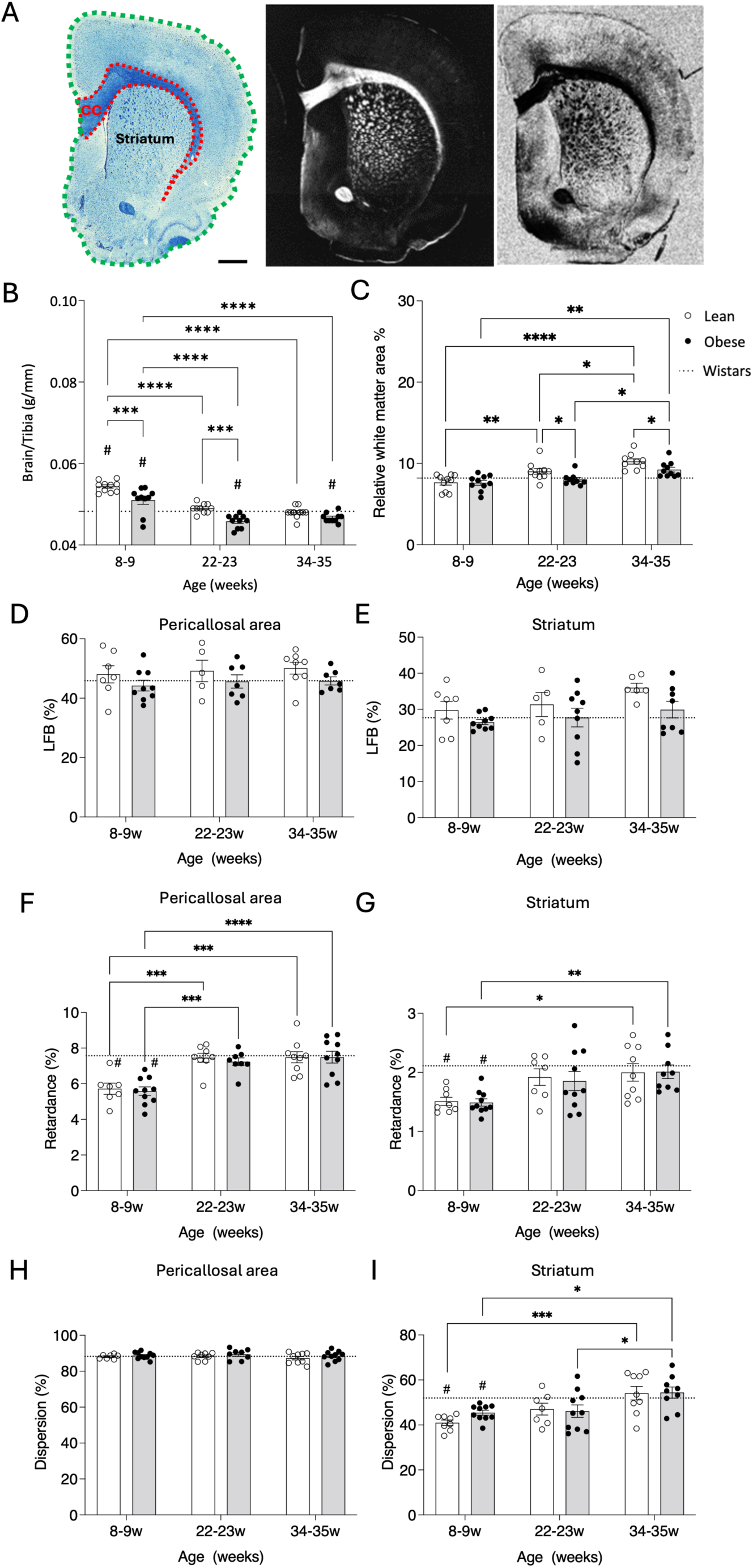
**Brain atrophy, white matter content and integrity in ZSF1 and Wistar rats**. (A) Representative image of a brain section stained with Luxol fast blue (left image) with delineation of the pericallosal area (red dotted line) and whole brain area (green dotted line); representative image depicting retardance (middle image) and dispersion (right image). Scale bar = 1 mm. (B) Brain weight to tibia length from Lean (open circles; white bars) and Obese (filled circles; grey bars) ZSF1 rats (2-way ANOVA: p_age_<0.0001, p_group_<0.0001, p_age x group_=0.13). (C) Relative proportion of the pericallosal area over whole brain tissue (2-way ANOVA: p_age_<0.0001, p_group_=0.03, p_age x group_=0.13). Relative Luxol Fast Blue intensity of the (D) pericallosal area (2-way ANOVA: p_age_=0,69, p_group_=0,04, p_age x group_=0.98) and (E) striatum area (2-way ANOVA: p_age_=0,09, p_group_=0,02, p_age x group_=0.79). Relative retardance of the (F) pericallosal area (2-way ANOVA: p_age_<0.0001, p_group_=0,62, p_age x group_=0.92) and (G) striatum area (2-way ANOVA: p_age_<0.001, p_group_=0,80, p_age x group_=0.95). Relative dispersion of the (H) pericallosal area (2-way ANOVA: p_age_=0,49, p_group_=0,17, p_age x group_=0.93) and (I) striatum area (2-way ANOVA: p_age_<0.0001, p_group_=0,051, p_age x group_=0.49). Sidak’s multiple comparisons test: *p<0.05, **p<0.01, ***p<0.001, ****p<0.0001. Single t-test: #p<0.05 vs (B) Wistar (black dotted line) = 0.04 ± 0.0008 g/mm, vs (C) Wistar = 8.19 ± 0.41 %. vs (D) Wistar = 45.8 ± 1.4 %. %. vs (E) Wistar = 27.6 ± 1.5 %. %. vs (F) Wistar = 7.5 ± 0.3 %. %. vs (G) Wistar = 2.1 ± 0.1 %. %. vs (H) Wistar = 88.2 ± 0.9 %. %. vs (I) Wistar = 51.9 ± 3.0 %. n=8-10/group.

### 4. Impaired neurovascular coupling in Obese ZSF1 rats

Cerebral blood flow was measured over the barrel cortex using laser speckle contrast imaging (LSCI) (**Figure 5A-C**). Baseline cortical cerebral blood flow and neurovascular coupling were assessed in ZSF rats at three different time points and in Wistar rats at 20-21w as a normotensive reference. Although baseline cerebral blood flow remained unchanged between Lean and Obese groups across all ages (**Figure 5D**), all ZSF1 animals had a significantly lower cerebral blood flow compared to Wistars at all time points (single t-test: p<0.05 vs Wistars). Moreover, blood flow declined significantly over time (p_age_=0.02), and a trend towards decreased baseline cerebral blood flow was observed in Obese rats at 32-33w vs 6-7w (p=0.0519) (**Figure 5D**). Neurovascular coupling, presented as the percentage increase in cerebral blood flow during whisker stimulation, was significantly decreased in Obese vs Lean at the second (p<0.001) and third (p<0.05) time point (**Figure 5E**). The impaired neurovascular coupling at 33 weeks of age was replicated in a separate study (**Figure S6**). In addition, cerebral blood flow increase to stimulation was significantly lower in ZSF1 rats at 6-7w, and in Obese animals at 20-21w and 32-33w compared to Wistar rats (single t-test: p<0.05 vs Wistars). Within Lean and Obese groups, the response to whisker stimulation was smaller at the first time point compared to 20-21w and 32-33w (p<0.01), which could be explained by an immature neurovascular coupling at 6-7w.

**Figure 5:**
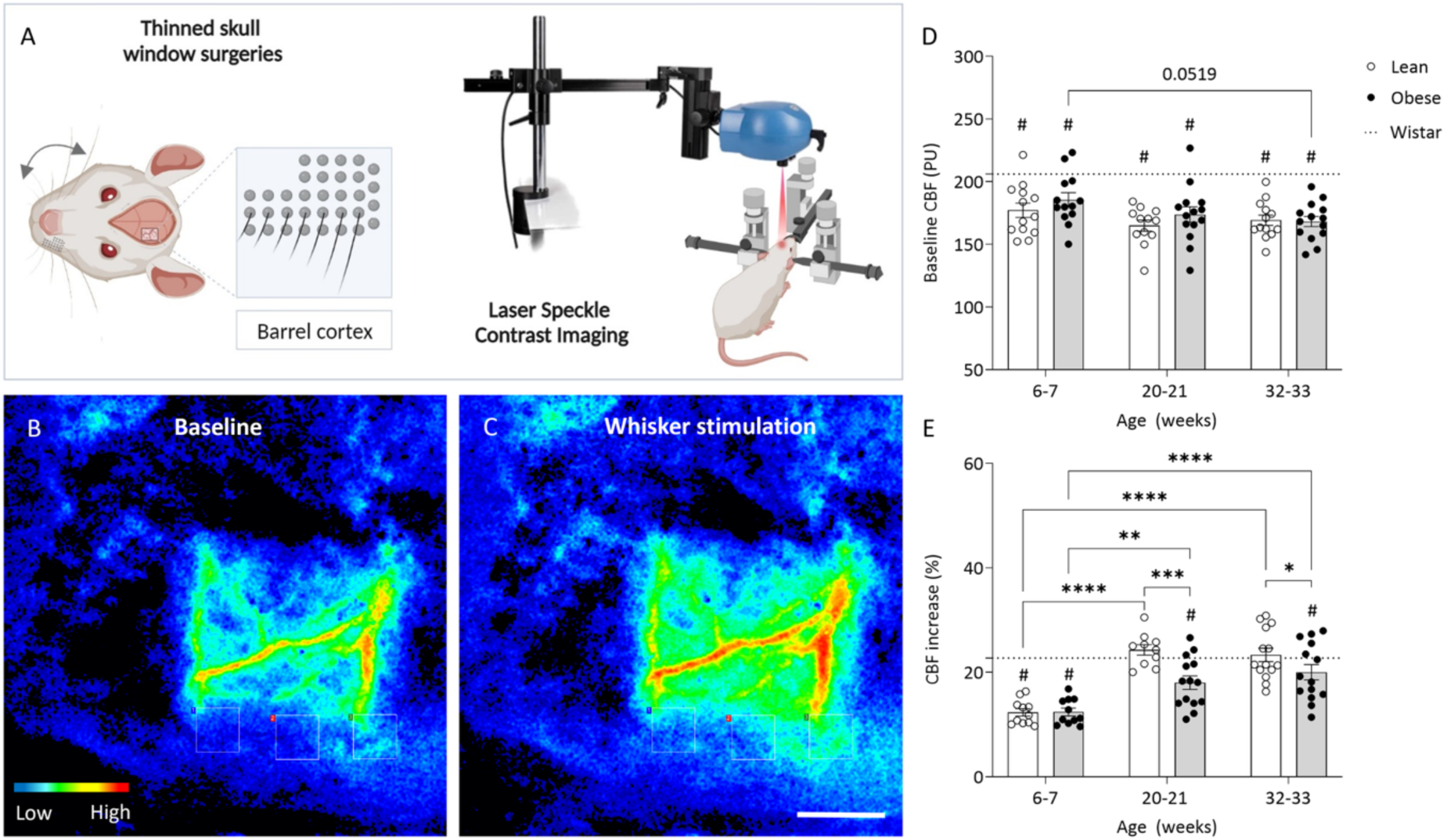
Cerebral blood flow and neurovascular coupling in ZSF1 and Wistar rats. (A) Schematic overview of thinned skull window surgeries and laser speckle contrast imaging over the barrel cortex area. (B-C) Representative images of cerebral blood flow (CBF) before and during whisker stimulation. (D) Baseline CBF (2-way ANOVA: p_age_=0.02, p_group_=0.20, p_age x group_=0.55) and (E) Neurovascular coupling (2-way ANOVA: p_age_<0.0001, p_group_=0.002, p_age x group_=0.04) of ZSF1 rats were assessed at three different time points and compared to normotensive Wistars rats (20-21w, dotted line). Sidak’s multiple comparisons test: *P<0.05, **p<0.01, ***P<0.001, ****P<0.0001. Single t-test: #p<0.05 vs (D) Wistar (20-21w) = 206 ± 10 PU and (E) Wistar = 23.42 ± 0.81 %. n=10-14/group. PU = perfusion units. Scale bar = 100 μm. Illustration in panel A was created using BioRender.com.

### 5. Obese ZSF1 rats showed increased cerebral vascular density and BBB leakage size

The microvascular network was assessed in the brain of ZSF1 rats at three different time points and in Wistar rats (22-23w). In addition to the group and interaction effect (p_group_=0.02, p_age x group_=0.03, **Figure 6A-E**), cortical vascular density was found to be higher in Obese vs Lean only at 34-35w (p<0.01). At this age, vascular density in the cortex was also significantly elevated compared to Wistar rats (single t-test: p<0.05 vs Wistars) and Obese rats at 22-23w (p<0.01). Although the density of branching points was similar between Lean and Obese groups, it was lower than in Wistar rats at 8-9w and 22-23w (single t-test: p<0.05 vs 349 ± 63 junctions/mm^2^, **Figure 6F**). In addition, the size of BBB leakages in the whole brain increased over time and was higher in the Obese group (p_age_=0.0026, p_group_=0.02). Whereas the number of BBB leakages **(Figure 6G-J**) per brain section was not different between Lean and Obese groups at any time point, the leakage size was significantly higher in Obese vs Lean at 22-23w (p<0.01). At this age, the size of BBB leakages within the Obese group was also higher than at 8-9w.

**Figure 6:**
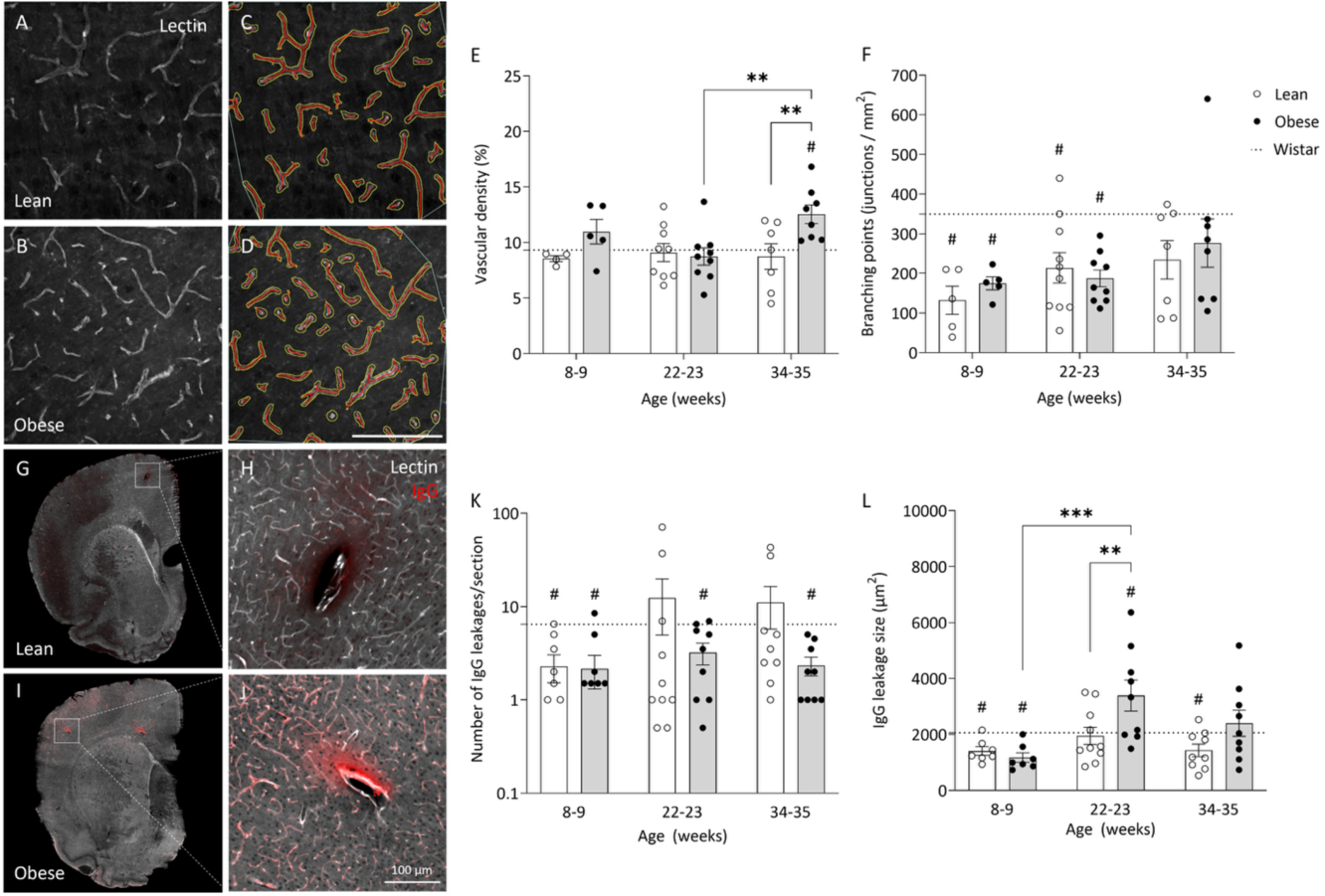
Cerebral microvasculature and BBB permeability in ZSF1 and Wistar rats. (A-B) Representative images (maximal projection) of lectin-stained brain sections in the deep cortical area prior to analysis, and (C-D) after analysis in Angiotool (Vessel width = yellow line; Vessel skeleton = red line; Branching points = blue dots). (E) Cortical vascular density (mixed effects: p_age_=0.12, p_group_=0.02, p_age x group_=0.03), and (F) Branching points (mixed effects: p_age_=0.10, p_group_=0.55, p_age x group_=0.61) in Lean (open circles; white bars) and Obese (filled circles; grey bars) ZSF1 rats. (G-J) Representative images of IgG leakages in brain sections from ZSF1 rats. (K) Number of IgG leakages (2-way ANOVA: p_age_=0.31, p_group_=0.06, p_age x group_=0.42), and (L) IgG leakage size (2-way ANOVA: p_age_=0.0026, p_group_=0.02, p_age x group_=0.09). Sidak’s multiple comparisons test: **p<0.01, ***p<0.001. Single t-test: #p<0.05 vs (E) Wistar (22-23w, dotted line) = 9.32 ± 0.63 %, (F) Wistar (22-23w, dotted line) = 349 ± 63 junctions/mm^2^, (K) Wistar (22-23w, dotted line) = 6.46 ± 1.44, and (F) Wistar (22-23w, dotted line) = 2057 ± 392.8 μm^2^. n=4-10/group.

### 6. Transcriptomic analysis of cortical microvessels

Isolated microvessels from the brain cortex of ZSF1 rats were characterized by immunofluorescent staining against markers for pericytes (PDGFR-β), microglia (Iba-1), astrocytic endfeet (aquaporin 4), and endothelial cells (tomato-lectin) (**Figure S7A-C**). Confocal imaging confirmed the presence of pericytes and astrocyte endfeet around the isolated cortical microvessels, whereas microglia could not be detected in the samples. The average diameter of the isolated vessels was 5.7 ± 1.2 μm (**Figure S7D**). Transcriptomic data revealed a total of 98, 254 and 487 differentially expressed genes (DEGs) in Obese vs Lean at 8-9w, 22-23w and 34-35w, respectively (p<0.05, absolute fold change>1.5; **Figure 7A-C**). Following Gene Ontology analysis, an overview of the main enriched biological processes was generated at each time point and further clustered into functional categories (22-23w in **Figure 7D**; 8-9w and 34-35w in **Figure S 8** and **Figure S9**). This analysis revealed a high regulation of processes related to angiogenesis and immune mechanisms in Obese vs Lean at the first time point. By 22-23w, these processes were still enriched, in addition to blood vessel formation and vasoreactivity. At the final time point, processes associated with blood vessel formation, vasoreactivity and vessel remodelling were significantly regulated.

**Figure 7:**
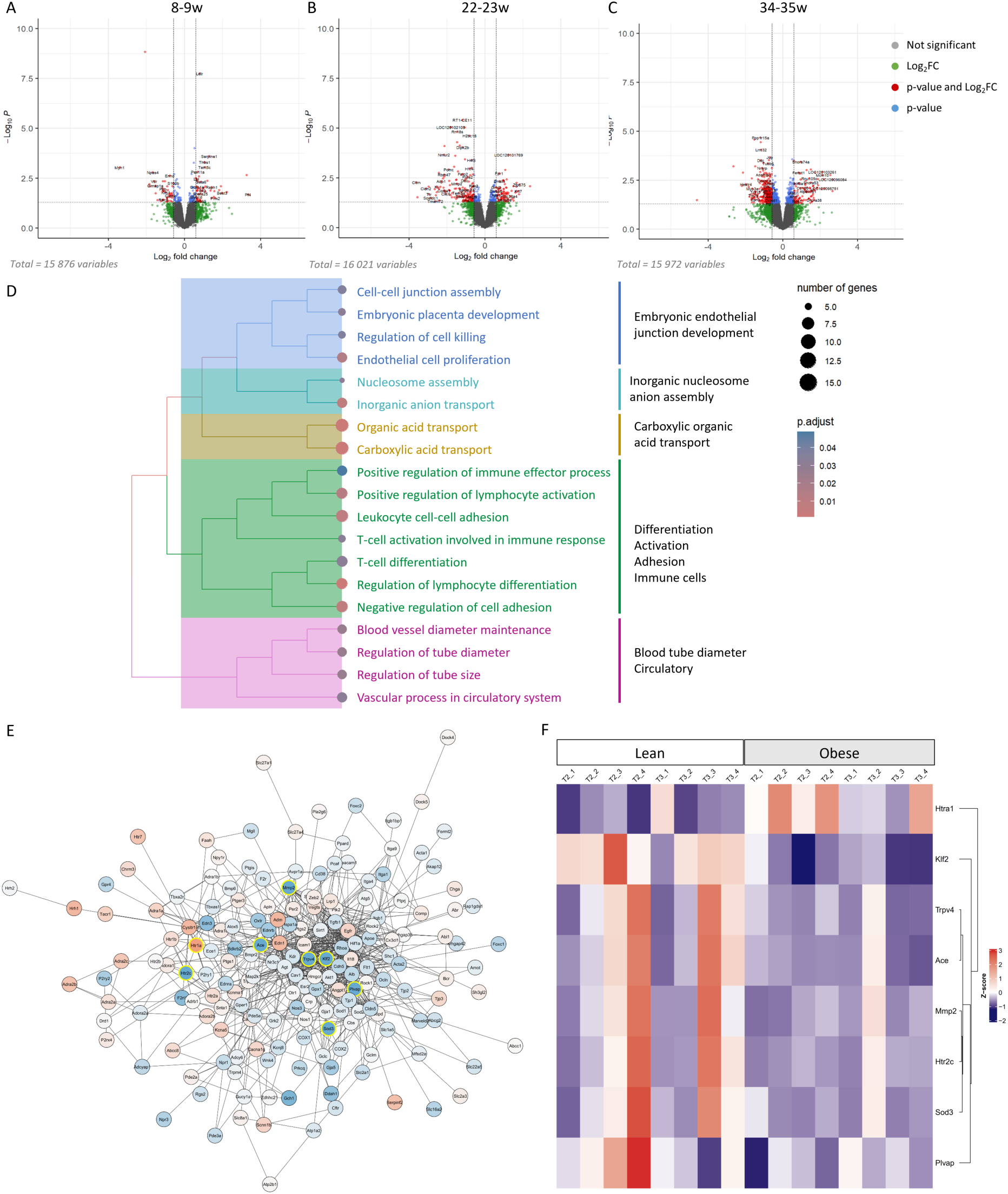
**Transcriptomic regulation within brain microvessels of ZSF1 rats**. (A-C) Volcano plots of all genes in the dataset, p-value<0.05 and fold change>1.5 (|log2FC|>0,58). The total number of differentially expressed genes (DEGs, Obese vs Lean) is 98 at 8-9w, 254 at 22-23w, and 487 at 34-35w.(D) Biological processes enriched within the dataset of 22-23w clustered into functional categories. (E) Protein-protein interaction network within the GO term GO:0003018 ‘vascular process in circulatory system’: DEGs (yellow circles) are represented with their direct connections. (F) Heatmap showing the expression profiles of DEGs within the GO term GO:0003018 at 22-23w and 34-35w: High-Temperature Requirement A Serine Peptidase 1 (Htra1), Kruppel-Like Factor 2 (Klf2), Transient Receptor Potential Vanilloid 4 (Trpv4), Angiotensin-Converting Enzym (Ace), Matrix Metallopeptidase 2 (Mmp2,) 5-Hydroxytryptamine (Serotonin) Receptor 2C (Htr2c), Superoxide Dismutase 3 (Sod3), and Plasmalemma Vesicle Associated Protein (Pvlap). n=4/group. Colour legend: red = upregulated, blue = downregulated.

Within these biological processes, Transient Receptor Potential Vanilloid 4 (*Trpv4*) and Krüppel-Like Factor 2 (*Klf2*), two genes that are involved in many of the enriched processes, were highly downregulated at both the second (p*_Klf2_*=0.00007, log2FC_Klf2_=-1.33; p*_Trpv4_*=0.02, log2FC=-1.18) and third (p*_Klf2_*=0.0004, log2FC*_Klf2_*=-1.04; p*_Trpv4_*=0.04, log2FC*_Trpv4_*=-1.21) time point in Obese vs Lean. Moreover, they were specifically regulated within the Obese groups but not in the Lean groups over time. Both genes are involved in processes related to ‘regulation of tube diameter’ (p_adj_=0.02), ‘blood vessel diameter maintenance’ (p_adj_=0.02), ‘regulation of tube size’ (p_adj_=0.02), and ‘vascular process in circulatory system’ (p_adj_=0.03) at the second time point, and the process ‘vasodilation’ (p_adj_=0.003) at the last time point. The protein-protein interaction network of all proteins associated with the overarching GO term GO:0003018 ‘vascular process in circulatory system’ is represented in **Figure 7E**.

The DEGs within this process include Requirement A Serine Peptidase 1 (Htra1), Klf2, Trpv4, Angiotensin-Converting Enzym (Ace), Matrix Metallopeptidase 2 (Mmp2,) 5-Hydroxytryptamine (Serotonin) Receptor 2C (Htr2c), Superoxide Dismutase 3 (Sod3), and Plasmalemma Vesicle Associated Protein (Pvlap), and their expression profile at 22-23w and 34-35w is represented in a heatmap in **Figure 7F**, with on average a lower expression level in Obese vs Lean, except Htra1 which is upregulated. Among these DEGs, *Trpv4* and *Klf2* are interacting with 32 and 17 other proteins, making them very central in this GO term.

### 7. Multiplex detection of *Klf2* and *Trpv4* expression in ZSF1 brains

In order to validate the downregulation of *Klf2* and *Trpv4* identified from cortical microvessels at 22-23w and 34-35w, a multiplex RNAscope was performed on paraffin-embedded brains from the same Lean and Obese animals used for the RNA sequencing (**Figure 8A-D, Figure S10**). The number of transcripts and total intensity of *Klf2* expression (**Figure 8E-F**) was significantly decreased in Obese vs Lean at 22-23w (p<0.001). The number of *Trpv4* transcripts was decreased (p<0.05) with a trend towards a lower signal intensity (p=0.06) in the Obese animals at this timepoint (**Figure 8G-H**). At the age of 34-35w, however, no significant differences were observed in the number or intensity of *Klf2* and *Trpv4* transcripts. In addition, two-way ANOVA indicates a group and interaction effect for both the number (p_group_=0.01, p_age x group_=0.0009) and intensity (p_group_=0.01, p_age x group_=0.0005) of *Klf2* transcripts, whereas only the interaction effect is observed in the number (p_age x group_=0.02) and intensity of *Trpv4* signal (p_group x age_=0.04). In summary, the downregulation of both targets could be confirmed at the second, but not at the third timepoint.

**Figure 8:**
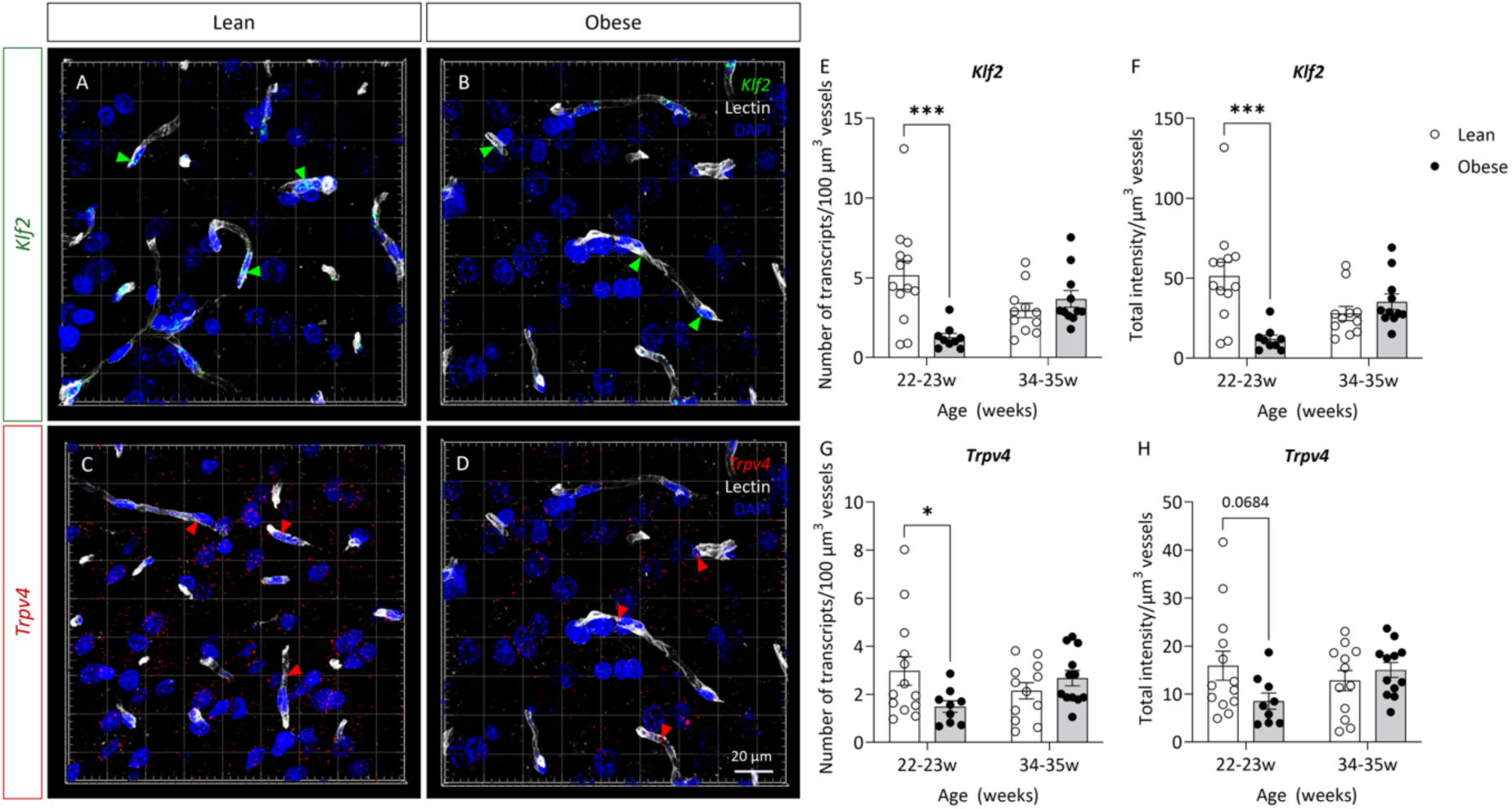
Multiplex detection of target probes Klf2 and Trpv4 in paraffin-embedded ZSF1 rat brains. (A-D) Representative images (maximal projection) of KLF2 (green) and TRPV4 (red) RNA transcripts inside lectin-stained vessels (grey) in the deep cortical area of Lean and Obese ZSF1 rat brains at 22-23 and 34-35 weeks of age. (E) The number of Klf2 transcripts per 100µm vessels (2-way ANOVA: p_age_=0.87, p_group_=0.01, p_age x group_=0.0009) and (F) total Klf2 signal intensity per µm^3^ vessels (2-way ANOVA: p_age_=0.99, p_group_0.01, p_age x group_=0.0005) in Lean (open circles; white bars) and Obese (filled circles; grey bars) ZSF1 rats; (I) Number of Trpv4 transcripts per 100µm vessels (2-way ANOVA: p_age_=0.67, p_group_=0.27, p_age x group_=0.02), (J) Total intensity of Trpv4 signal per µm^3^ vessels (2-way ANOVA: p_group_=0.27, p_age_=0.46, p_group x age_=0.04). Sidak’s multiple comparisons test: *p<0.05, ***p<0.001. n=10-13 FOVs/group.

## DISCUSSION

Alongside confirming the development of the comorbidities and HFpEF in this model, we demonstrated progressive cognitive impairment in Obese rats starting from 20 weeks of age. Obese animals had short-term memory impairment, altered spatial learning, and impaired long-term memory, while working memory and reversal learning remained similar to Lean rats. As expected, mild cognitive decline was also observed in Lean hypertensive animals at the final time point, as suggested by the impaired working memory and the smaller difference in performance between Lean and Obese during the Barnes Maze task. The overall cognitive dysfunction was furthermore associated with brain atrophy and decreased myelin content in Obese ZSF1 rats.

In addition to cognitive decline, the longitudinal study of ZSF1 rats provided more insight into the chronic impact of comorbidities on cerebrovascular reactivity. Endothelial dysfunction has been recognized as an important hallmark of cSVD, as confirmed in a systematic review and meta-analysis of 29 studies, who reported a reduced neurovascular coupling in different forms of cSVD (34). Our findings indicated that impaired neurovascular coupling is present in Obese ZSF1 rats starting from 22-23w, and this was associated with the development of cognitive impairment. The response to whisker stimulation was also found to be lower at 6-7w in both ZSF1 groups compared to later time points and Wistars, possibly due to an immature vascular reactivity at this age (35, 36). Although no significant difference in baseline cerebral blood flow was detected at any time point, it did decrease over time. Given the limited imaging depth of LSCI, it is also possible that perfusion in deeper regions of the brain is affected to a greater extent.

Together with the development of cognitive impairment and neurovascular dysfunction, Obese ZSF1 rats exhibited brain atrophy, a common feature in cSVD. By applying a multimodal assessment of white matter area, myelin content, density, and integrity, we were able to detect mild, yet meaningful microstructural defects in Obese ZSF1 rats. In addition to reduced brain volume, Obese animals showed subtle but consistent reductions in white matter area, in myelin content and early signs of compromised myelin quality within the striatum. Age-related increases in myelin density were evident in both Lean and Obese groups, likely reflecting ongoing maturation at this stage(36–38). Altogether, these data support the concept that comorbid metabolic stress and neurovascular uncoupling begin to drive structural white matter vulnerability early in the disease course.

Considering the cognitive, neurovascular, and white-matter deficits identified in this model, we next assessed whether comorbidities extend their impact to the structure and permeability of the cerebral microvasculature. Immunohistochemical analysis showed an increase in both BBB leakage size and microvessel density at the second and third time point, respectively. Some controversy exists on the vascular density in cSVD; various papers have reported decreased vascular density while others observed an increase, similar to our findings. It is well established that high blood pressure, one of the risk factors for cSVD, is associated with capillary rarefaction, as observed by Suzuki et al. in renal hypertensive rats (39). On the other hand, Liu et al. confirmed enhanced neovascularization as well as microvascular hypoperfusion in adult leptin-receptor deficient mice (40). In addition, vascular density and permeability were elevated in obese mice fed with a high-fat diet (41). Our data suggest that, despite the increased number of microvessels, these new vessels are more unstable and dysfunctional, and therefore unable to properly respond to the needs of the brain’s parenchyma. We propose that the observed enhanced vascularization at the latest time point serves as a compensatory mechanism for the chronic hypoperfusion and impaired cerebrovascular reactivity in Obese ZSF1 rats. The resulting increase in permeability may further initiate or exacerbate an inflammatory cascade.

Our study aimed at examining the molecular processes central to the development of VCI in this HFpEF model by studying the transcriptomic regulations within cortical brain microvessels at three different time points. The Gene Ontology analysis revealed a clear evolution of regulated processes over time. At the first time point, many processes were related to immune mechanisms, which were still detected at the second time point. Subsequently, processes related to angiogenesis and vasodilation became highly enriched, highlighting the functional changes in the vasculature. By 34-35w, the enrichment of processes linked to angiogenesis, vasoreactivity, and vessel remodelling emphasize ongoing structural changes. These data align with the observed impaired neurovascular coupling starting at 22-23w, as well as the increased vessel density at the final stage.

We further validated *Trpv4* and *Klf2* as important targets in processes related to vascular reactivity. Notably, *Trpv4* and *Klf2* downregulation were specific to the Obese group; a decrease in expression was observed at both the second and third time point, while this reduced expression was not detected in Lean rats. The Ca^2+^-permeable TRPV4 channel is known to play a crucial role in regulating vascular tone, angiogenesis, BBB integrity, and neuro-inflammation (42–44). A decline in NO production was observed when *Trpv4* was downregulated, resulting in impaired vasodilation (43). In addition, TRPV4-mediated dilation has been associated with cognitive function and cerebrovascular reactivity (45, 46). Secondly, *Klf2* is involved in anti-inflammatory, anti-angiogenic and anti-oxidant activities, and is known to trigger eNOS expression in endothelial cells, thereby inducing vasodilatory effects (47). Reduced expression of *Klf2* was found in the brain of Alzheimer patients, and upregulation of endothelial *Klf2* has been shown to attenuate Aβ-induced cytotoxicity and improve mitochondrial function (48). We were able to validate the downregulation of *Trpv4* and *Klf2* at 22-23w but not at 34-35w. This discrepancy might be explained by the difference in the material used between the two approaches (isolated microvessels vs endothelial cells). Since the transcriptomic analysis was performed on bulk RNA extracts from brain microvessels, comprising endothelial cells, pericytes and astrocyte endfeet, we cannot elaborate on the respective cell type contribution. Nevertheless, it is well described that *Trpv4* and *Klf2* have a more prominent function in endothelial cells (49, 50).

Beyond the diagnostic challenges, current treatment strategies primarily focus on slowing the progression of comorbidities (e.g. hypertension, type II diabetes), but no specific targeted therapy exists for HFpEF nor VCI. Taken together, our results indicate for the first time that VCI is present during HFpEF in ZSF1 rats. The presence of comorbidities and spontaneous development of both cardiac and cerebral pathologies make this model unique and highly translational to investigate mechanisms underlying cSVD. This study demonstrates the downregulation of *Trpv4* and *Klf2,* which was associated with the onset of neurovascular uncoupling, as well as increased BBB permeability and vascular density. Given their mechanosensitive and vasoprotective properties, *Trpv4* and *Klf2* could be promising therapeutic targets for cerebrovascular dysfunction. Although two clinical trials reported the safety of the TRPV4 antagonist GSK2798745 in healthy volunteers and patients with heart failure, no clinical trials have investigated the use of TRPV4 agonists/antagonists in the context of neurological dysfunction (51, 52). Due to the lack of specific treatments for cSVD and VCI, we suggest to further explore the potential of these targets in other preclinical models, human tissue and clinical trials.

## Declarations

### Ethics approval and consent to participate

All experiments and procedures within this study were conducted in accordance with institutional guidelines and approved by the Ethical Committees for Animal Experiments of Maastricht University (AVD10700202010326).

### Data availability of data and materials

Data are available upon reasonable request.

## Funding

This research was supported by the European Union’s Horizon 2020 project ‘CRUCIAL’ (grant number 848109) and by Horizon 2020 research and innovation programme under the Marie Skłodowska-Curie (grant agreement No 954798).

### Authors’ contributions

Conceptualization and study design: S.L., S.F.; Acquisition of research funding: S.F., E.J., R.v.O.; Performed in vivo and ex vivo experiments and data analysis: S.L.,L.v.T., I.P., P.L., N.B., D.M., D.H., S.S., E.W., H.S., F.A., M.W.; Performed the RNA sequencing study: M.v.H., R.K.; Performed and supported bioinformatic analyses: S.L., S.F., F.C., M.K.; Supervision and guidance: S.F., M.W., A.J.K., E.J., R.v.O.; All the authors contributed to the writing and reviewing of the paper and approved the final version.

## Acknowledgements

Alessandra Pasut and Mieke Dewerchin, Laboratory of Angiogenesis and Vascular Metabolism, Department of Oncology and Leuven Cancer Institute (LKI), KU Leuven, VIB Center for Cancer Biology, VIB, Leuven, Belgium. All partners from the Horizon 2020 project ‘CRUCIAL’.

---

**Figure S1.**
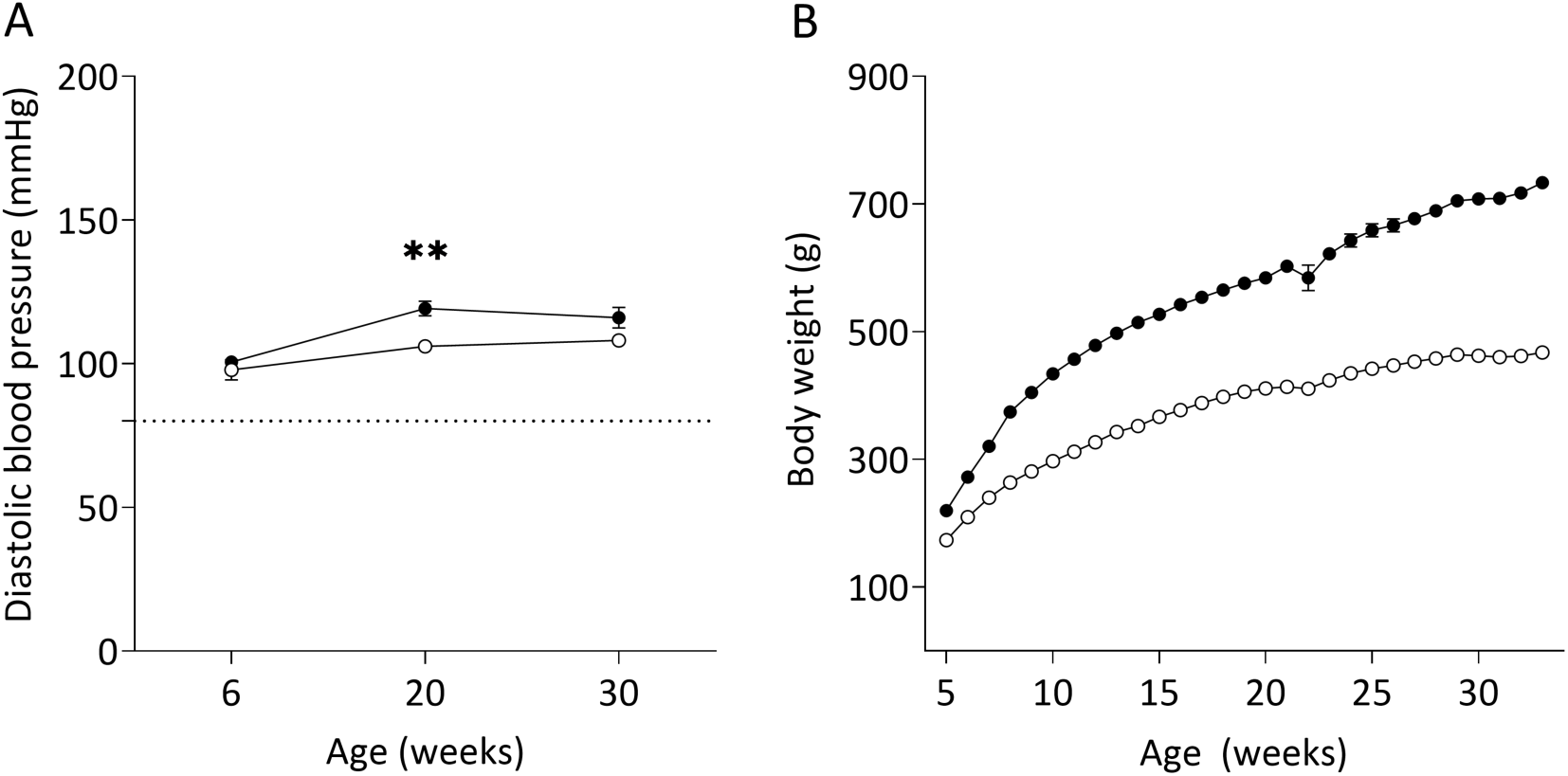

**Figure S2.**
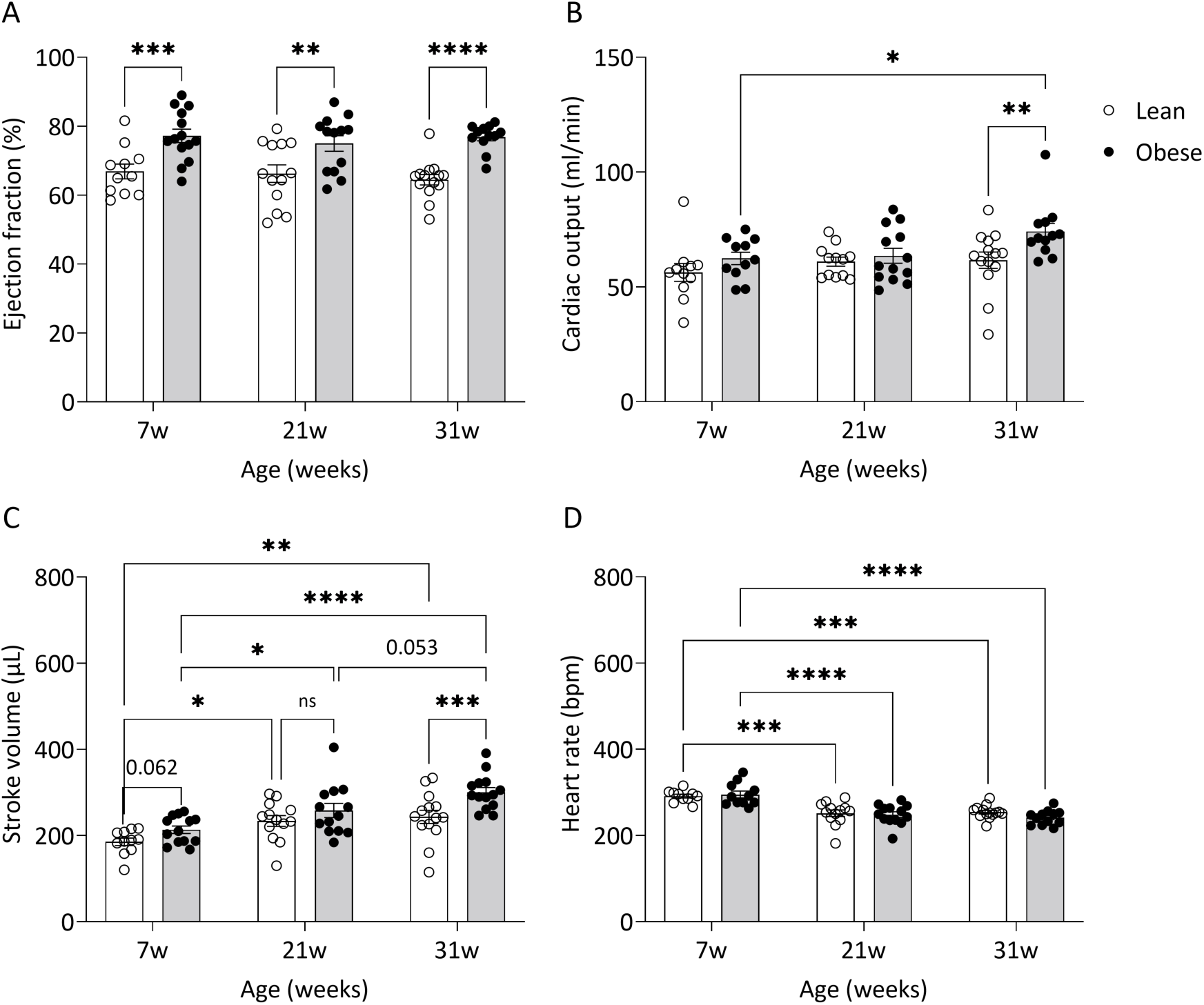

**Figure S3.**
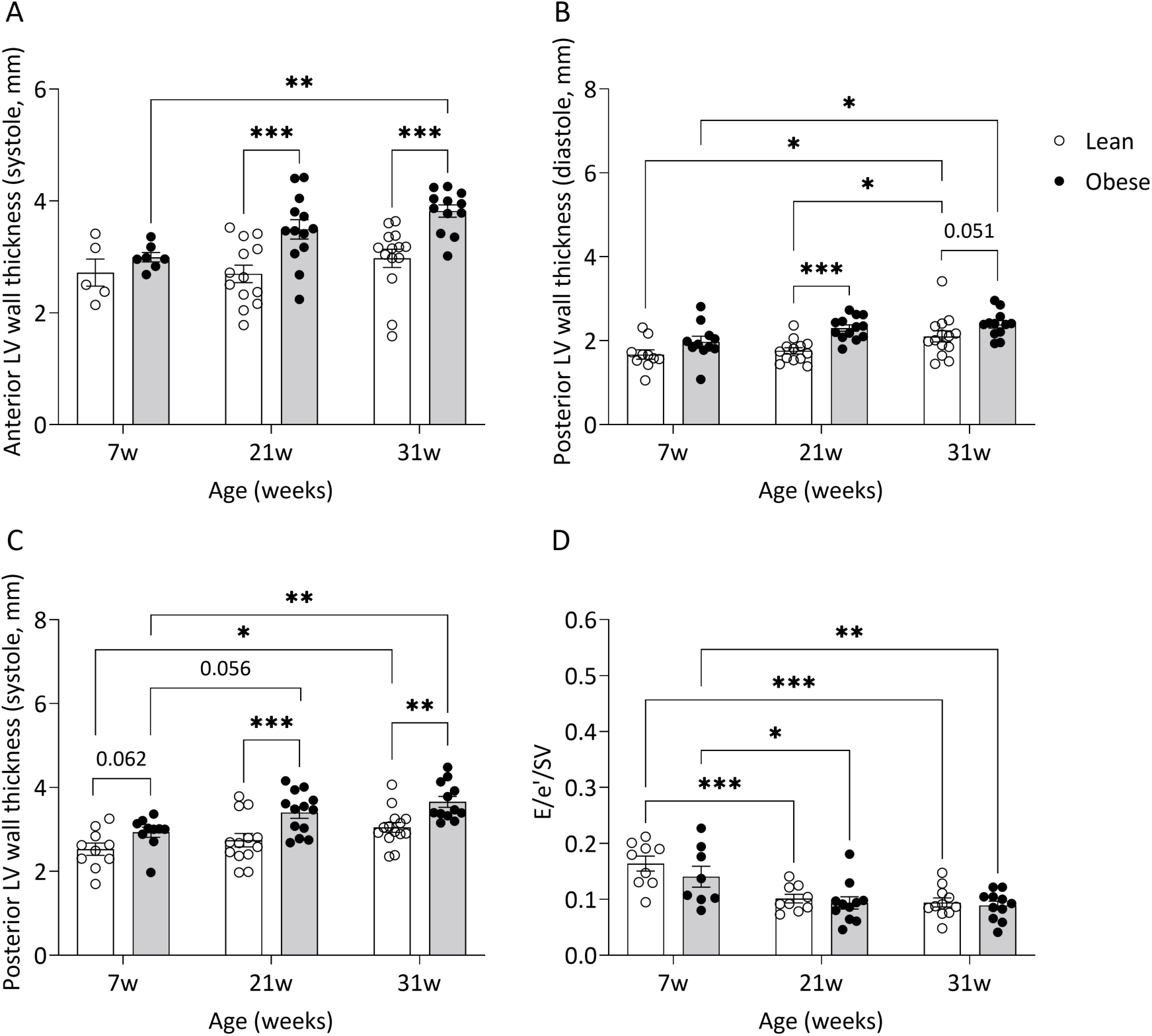

**Figure S4.**
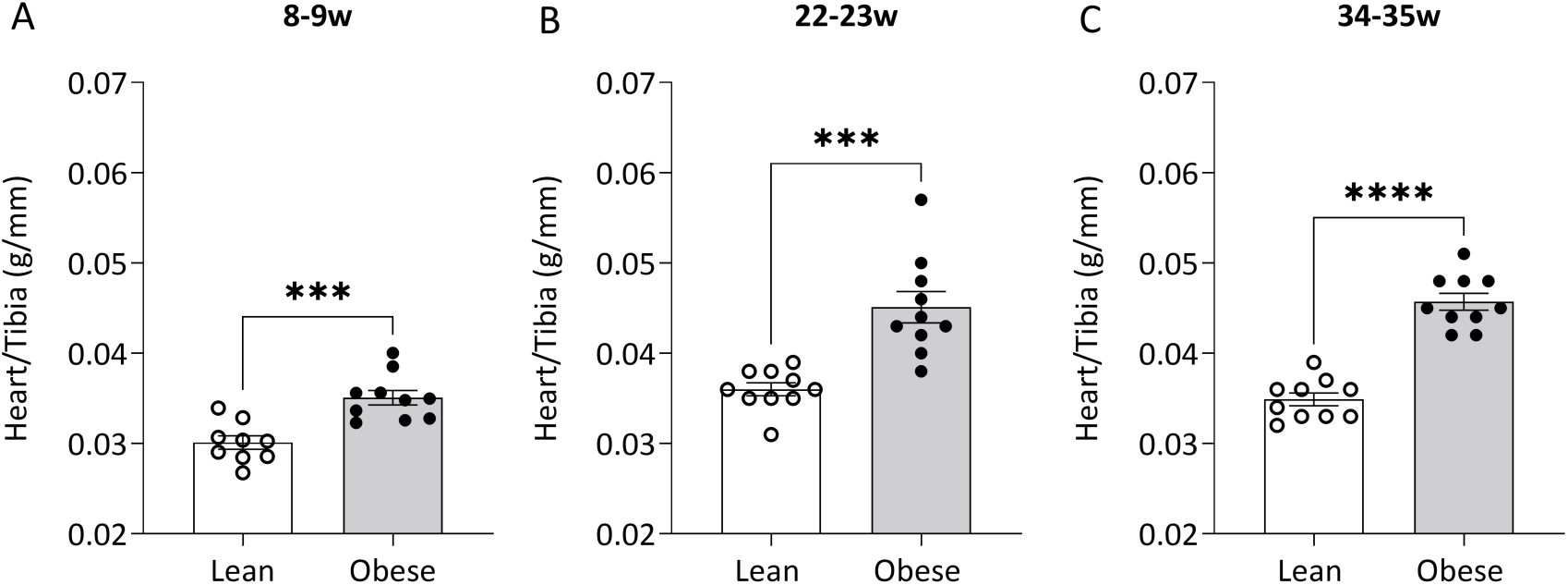

**Figure S5.**
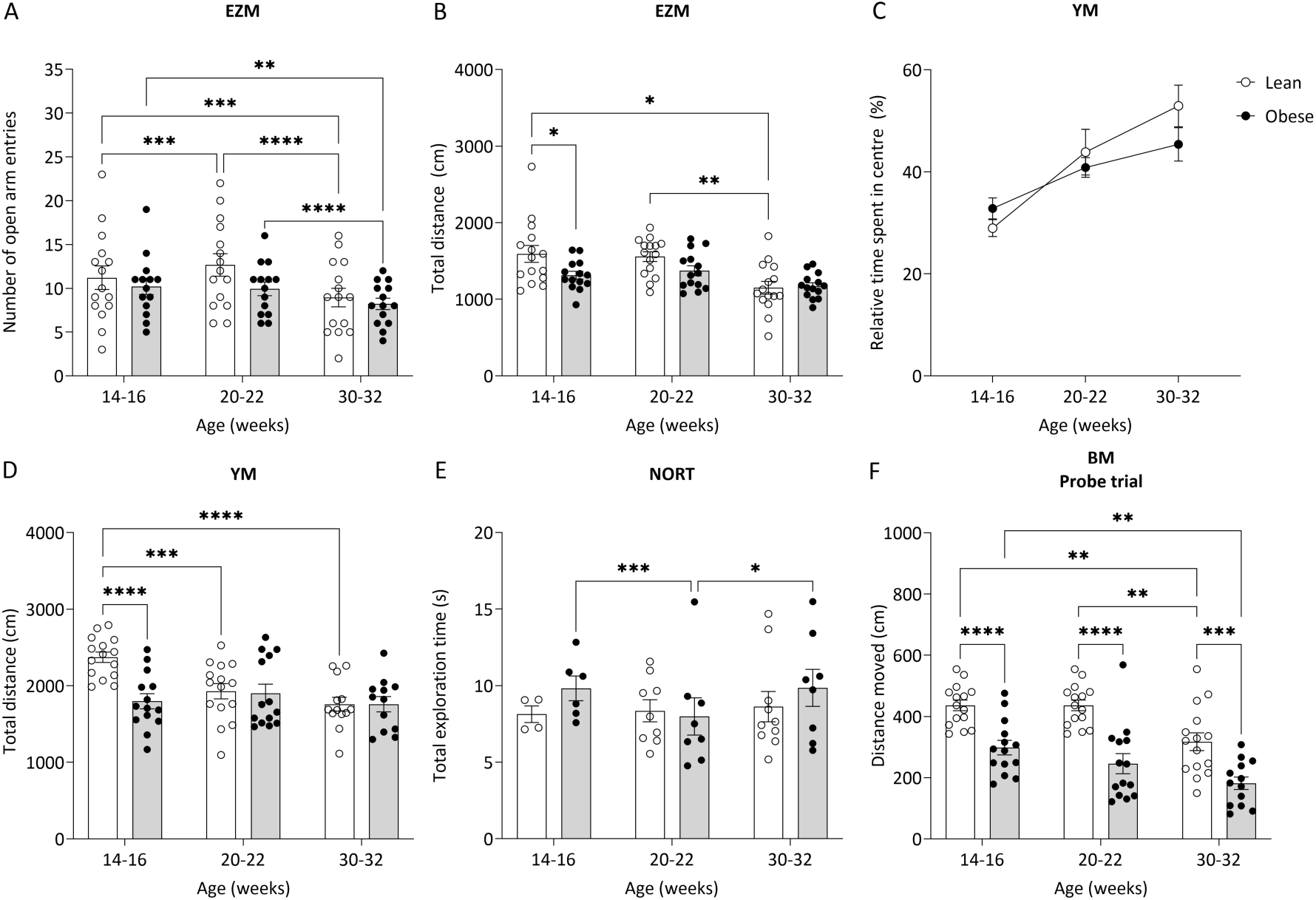

**Figure S6.**
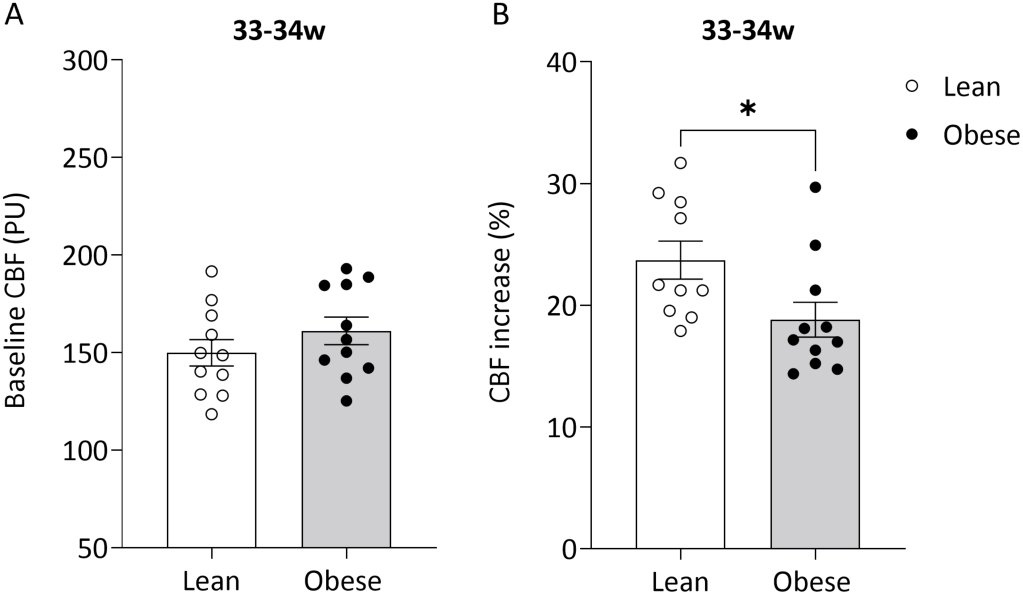

**Figure S7.**
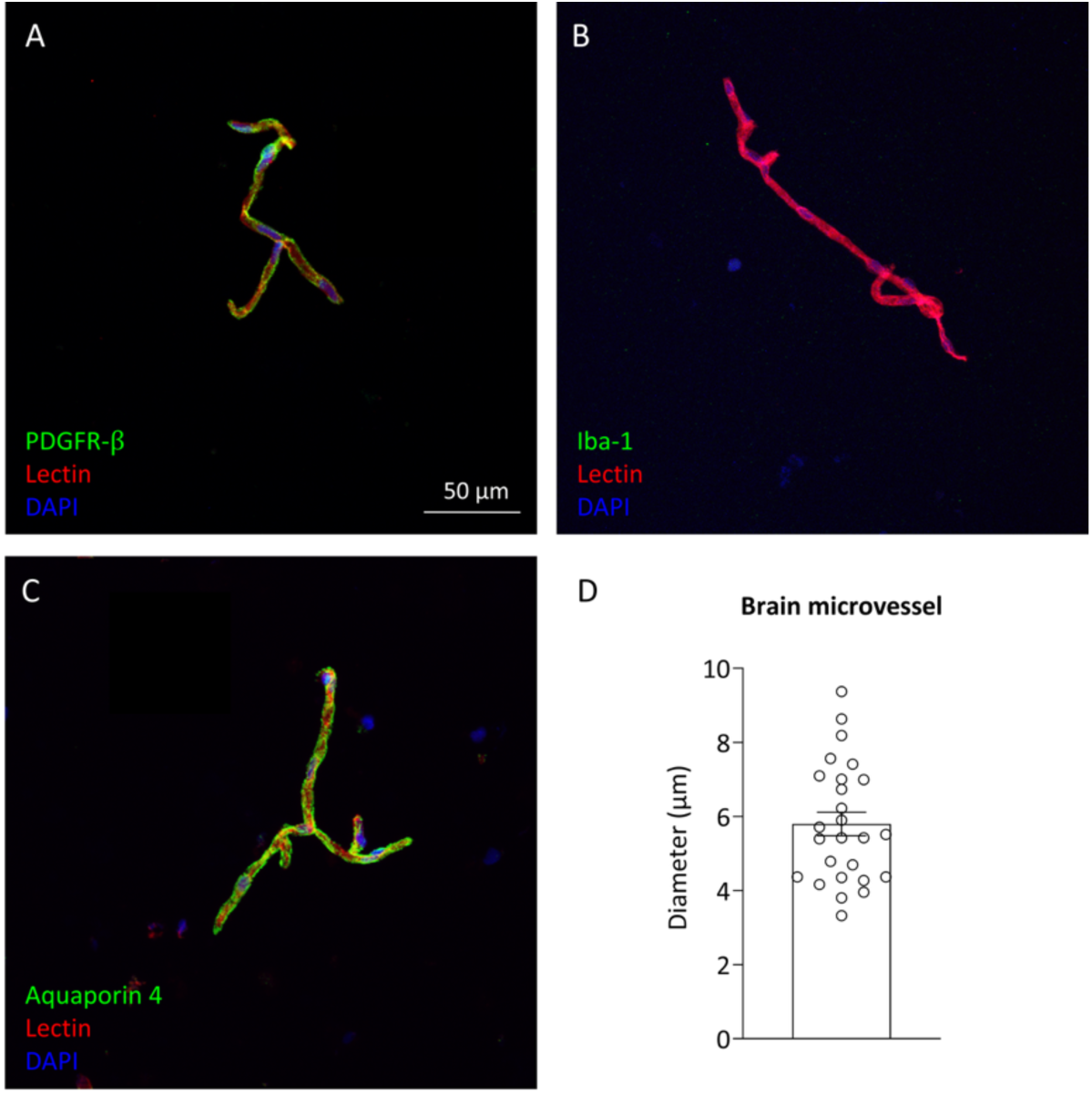

**Figure S8.**
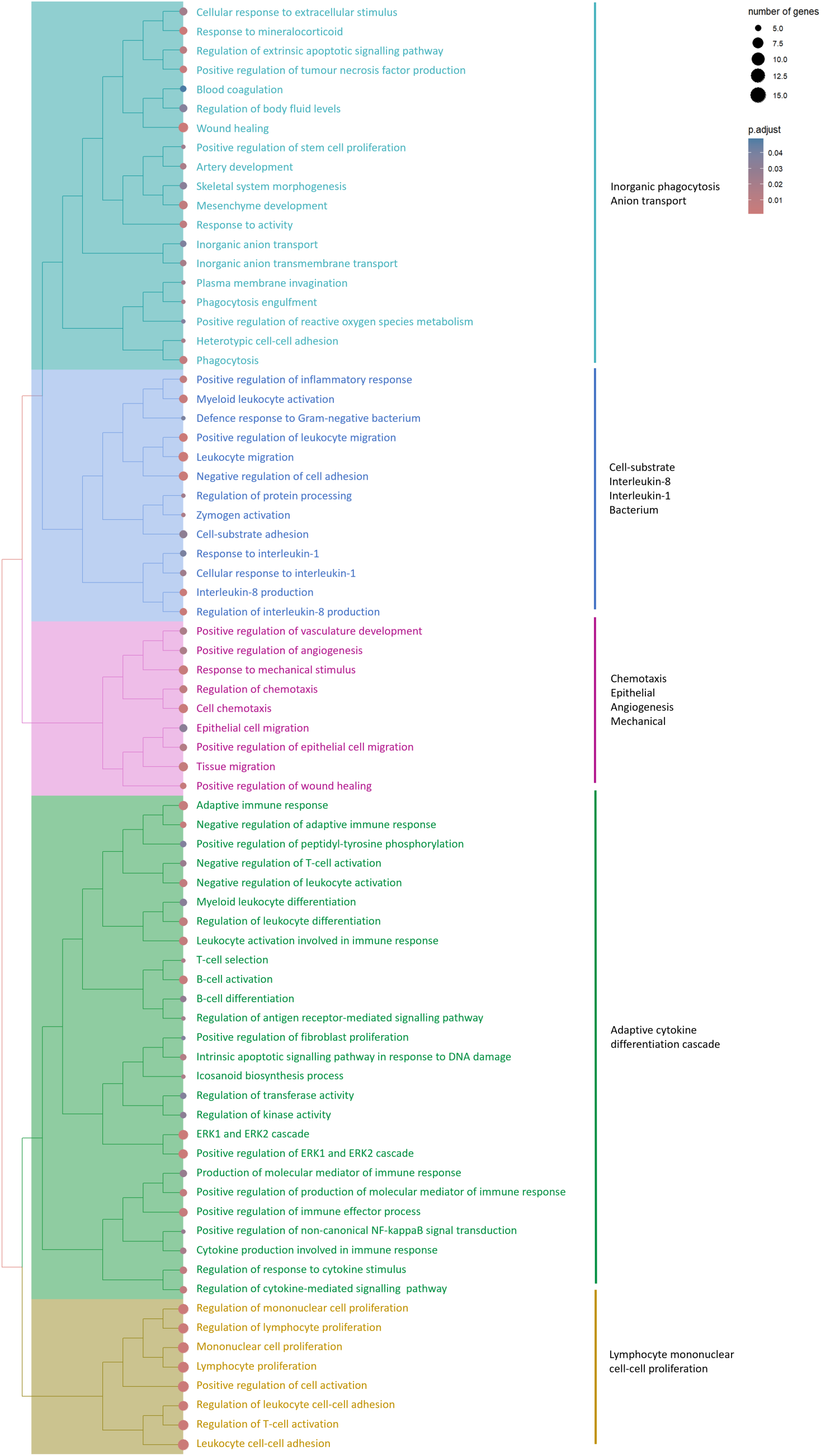

**Figure S9.**
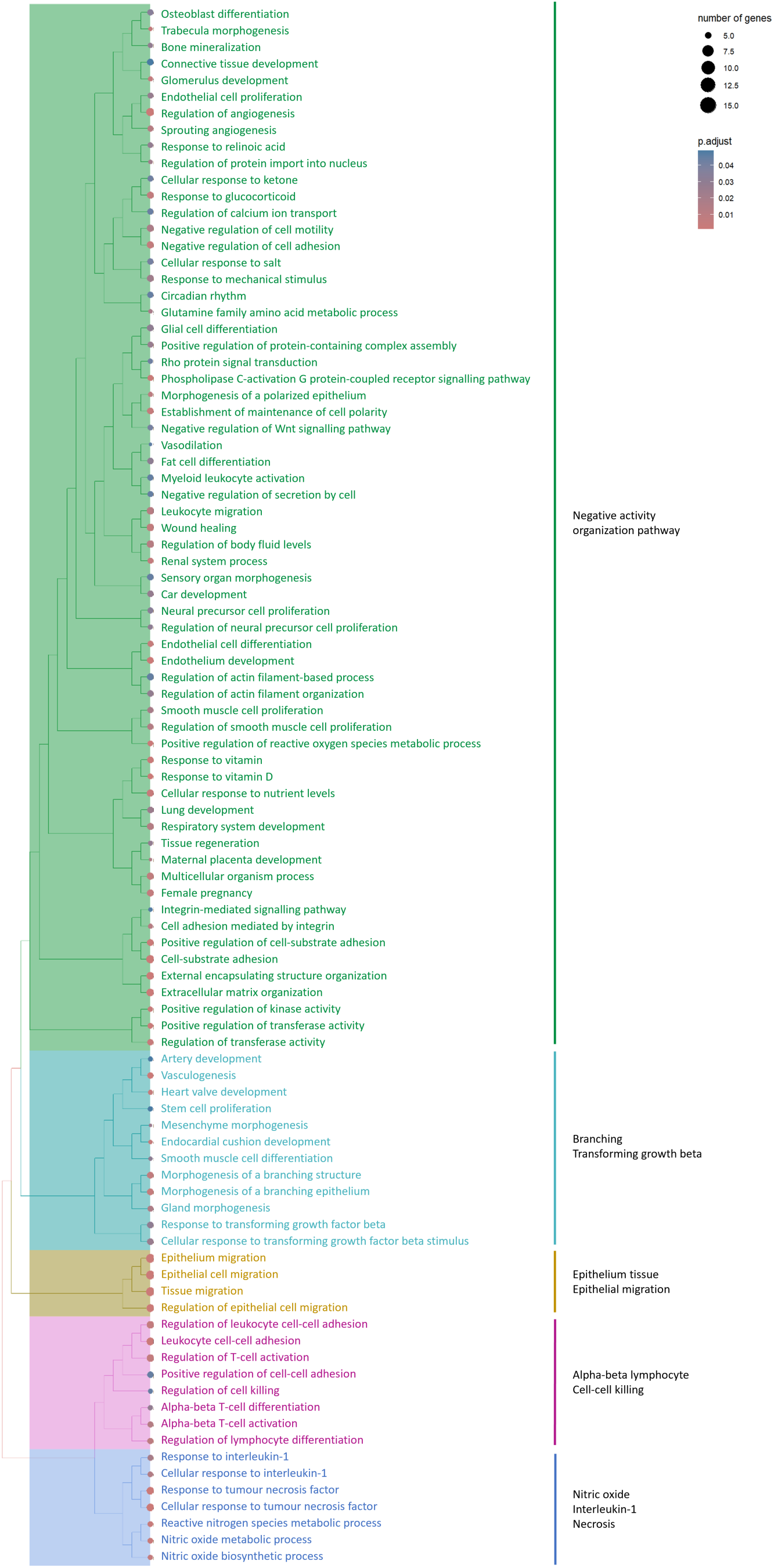

**Figure S10.**
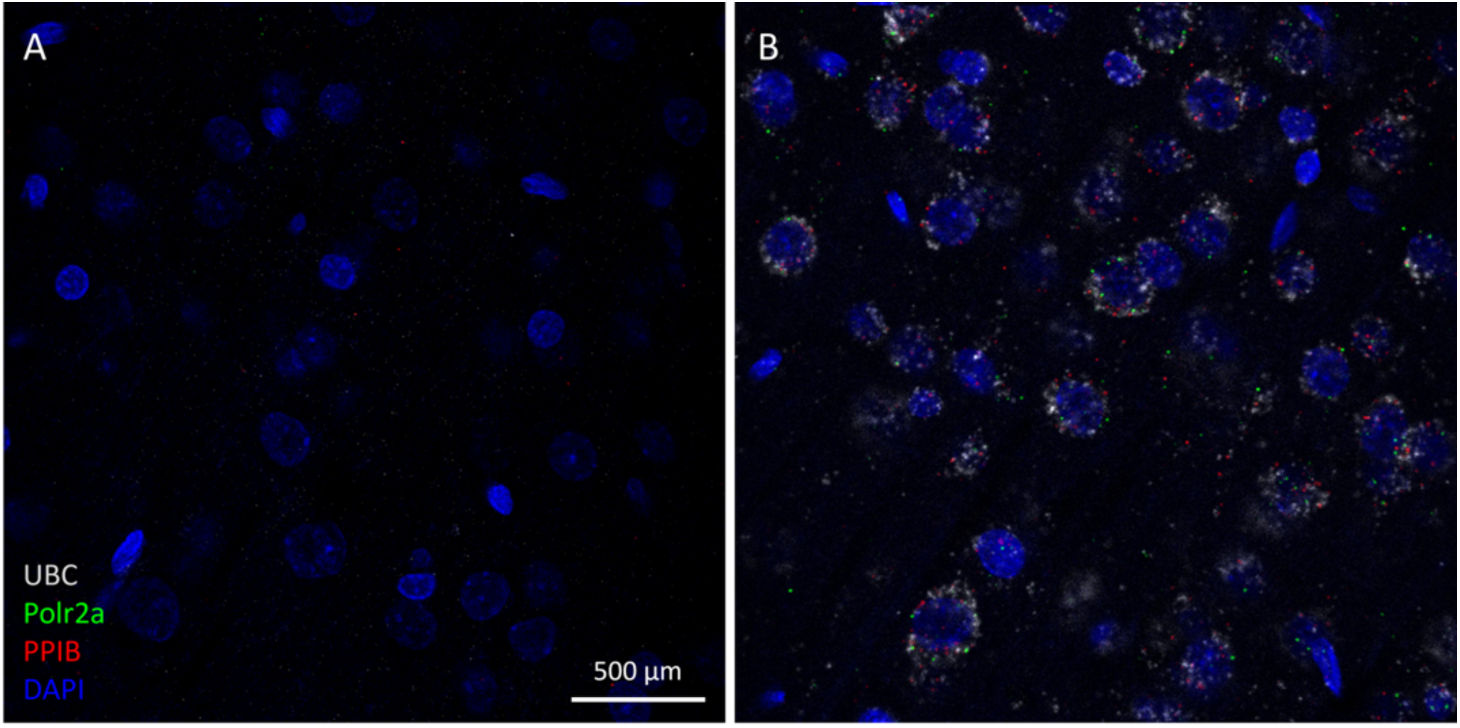

